# NF-κB signaling directs a program of transient amplifications at innate immune response genes

**DOI:** 10.1101/2025.03.11.641929

**Authors:** Michael P. Ludwig, Jason R. Wilson, Matthew D. Galbraith, Nirajan Bhandari, Lauren N. Dunn, Joshua C. Black, Kelly D. Sullivan

## Abstract

The cellular response to pathogens involves an intricate response directed by key innate immune signaling pathways which is characterized by cell-to-cell heterogeneity. How this heterogeneity is established and regulated remains unclear. We describe a program of transient site-specific gains (TSSG) producing extrachromosomal DNA (ecDNA) of immune-related genes in response to innate immune signaling. Activation of NF-κB drives TSSG of the interferon receptor gene cluster through inducible recruitment of the transcription factor RelA and the pre-replication complex member MCM2 to an epigenetically regulated TSSG control element. Targeted recruitment of RelA or p300 are sufficient to induce TSSG formation. RelA and MCM2 specify a program of TSSG for at least six and as many as 179 regions enriched in innate immune response genes. Identification of this program reveals regulated production of ecDNA as a mechanism of heterogeneity in the host response.

## INTRODUCTION

Innate immunity is the first line of defense against pathogens, active in every cell in the host, and requires no permanent genetic changes. This evolutionarily ancient response to pathogens results in rapid changes in cell signaling to inform both the infected cell and surrounding cells of potential danger. In higher eukaryotes, a complex network of molecular pattern recognition receptors (PRRs) senses pathogen- and damage-associated molecular patterns (PAMPs and DAMPs), including exogenous nucleic acids from viral or bacterial infection, and triggers a robust signaling cascade resulting in activation of numerous transcriptional networks.^1-4^ This network is controlled by transcription factors including nuclear factor kappa-light-chain-enhancer of activated B cells (NF-κB) and interferon (IFN) regulatory factors 3 and 7 (IRF3 and IRF7), among others.^3,4^ Execution of this transcriptional program results in production and release of interferons and cytokines, signaling infection and directing cell autonomous functions to respond to and eliminate pathogens.^5-7^ These receptors, signaling pathways, and transcriptional responses, have been extensively studied in both immune and non-immune cell types resulting in key breakthroughs in our understanding of infectious disease, but also leading to new waves of therapeutics in cancer biology and beyond.^8-11^

It has become clear that many developmental, signaling, and stress response pathways exhibit stochastic gene expression at the cellular level. This transcriptional heterogeneity in a population of cells has emerged as key contributor to overall cellular responses including innate immune signaling.^12,13^ The production of many cytokines, including IFNs, occurs in only a fraction of cells in a population in response to activation of innate immune signaling.^14-19^ For example, single cell RNA sequencing (scRNA-seq) has demonstrated that fewer than 25% of fibroblasts or mononuclear cells induce transcription of cytokines upon stimulation.^15^ Additionally, not all cells produce the same cytokines in response to the same innate immune stimulus. However, it remains unclear how this heterogeneity in innate immune response is achieved and regulated.

Extrachromosomal DNA (ecDNA) have emerged as a key contributor to genetic and phenotypic heterogeneity in tumors.^20-26^ However, the presence of ecDNA in normal healthy cells including muscle, sperm, and immune cells, as well as detection in S. cerevisiae suggests an importance in normal biological functions.^27-29^ (ecDNA) describes DNA elements that reside outside of canonical chromosomes. Since initial reports in the 1960s, it has emerged that ecDNA can exist as linear or circular (extrachromosomal circular DNA; eccDNA) fragments.^30-37^ Unlike chromosomal DNA, which is reliably segregated and inherited in each cell division, ecDNAs are randomly distributed and thus a key feature of cell-to-cell heterogeneity.^38,39^ While ecDNA have been observed in many organisms, including humans, its prevalence, origins, and functional significance have remained largely unexplored until recently.^37^ The biological functions of ecDNA have largely been investigated in cancer where their presence has been correlated with genomic instability, amplification of oncogenes, and drug resistance, thereby contributing to tumor heterogeneity and therapeutic challenges.^21,37,40-42^ Moreover, ecDNA have been linked to the rewiring of cellular signaling networks, altered gene expression patterns, and aberrant chromatin organization, potentially influencing cellular phenotypes, drug sensitivity, and disease outcomes.^21,37,40-43^ Despite the growing recognition of the importance of ecDNA, numerous questions regarding the biogenesis, replication, and regulation remain unanswered.

Understanding the mechanisms underlying ecDNA biogenesis is essential to deciphering the functional significance of ecDNA and uncovering the therapeutic potential. Though small ecDNA in some cancer cells correlate with microdeletions found in those cells, the presence of larger ecDNA in normal cells is not consistent with the generation of ecDNA from large scale deletions.^44,45^ These data suggest that a mechanism to amplify genomic regions could be part of ecDNA biogenesis.^25,31,37,46^ One mechanism for targeted amplification for ecDNA production is DNA rereplication.

In eukaryotic organisms, replication proceeds through origin specification, licensing, firing and ultimately, DNA synthesis.^47,48^ Exquisite control mechanisms have evolved to control DNA origin of replication specification, licensing, and firing, ensuring faithful duplication of the genome.^47,48^ However, it has become apparent that under specific developmental or environmental conditions, targeted DNA rereplication transiently increases gene dosage.^49^

In response to stress, cells amplify specific genomic loci through targeted DNA rereplication.^49-54^ These Transient Site-Specific Gene amplifications (TSSGs), which are a type of ecDNA, are an evolutionarily conserved response to environmental stress.^50,55^ TSSGs are targeted DNA rereplication events that occur during S phase and result in a transient increase in gene dosage through ecDNA that is cleared prior to nuclear envelop breakdown in G2 (**Figure 1A**).^50,51^ This transient increase in gene copy number is not dependent on changes in chromatin compaction during cell cycle but requires an actively cycling cell in S phase.^49-54^ Transiently increasing gene dosage and transcript levels may enable cells to respond to stress conditions without the need to rewire entire transcriptional networks and contribute to cell to cell heterogeneity.

**Figure 1.**
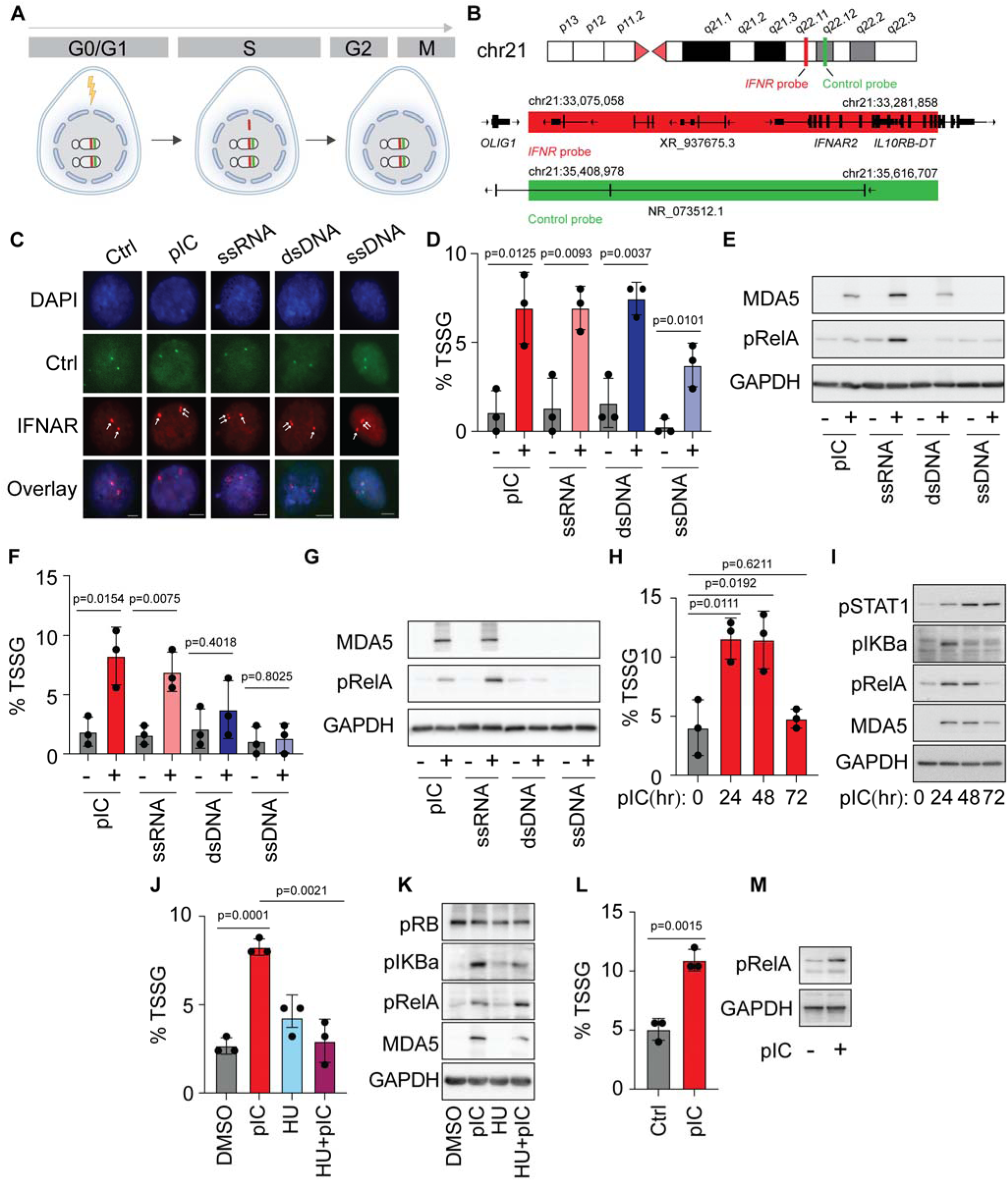
The interferon receptor locus on chr21 undergoes TSSG in response to innate immune activation. (**A**) Schematic of stimulus-induced production of TSSGs during S-phase which are lost by G2/M. (**B**) Schematic of interferon receptor (*IFNR*) locus (red) and control locus (green) fluorescence *in situ* hybridization (FISH) probe locations on human Chr21. (**C**) Representative images of *IFNR* (red, arrows) and control (green) foci from fibroblasts treated as indicated. Scale bars: 5 μM. (**D, F**) Percentage of cells undergoing *IFNR* TSSG in fibroblasts (D) and retinal pigment epithelial cells (RPE, E) transfected with the indicated nucleic acids (pIC 31.25 ng/mL, ssRNA 62.5 ng/mL, dsDNA 250 ng/mL, ssDNA 500 ng/mL) for 24 hours (n = 3). TSSG was scored as the percentage of nuclei with more *IFNR* foci (red) than control foci (green). (**E, G**) Immunoblot analysis of immune response activation from samples in D and F. GAPDH was used as a loading control. (**H**) Percentage of fibroblast cells undergoing *IFNR* TSSG transfected with pIC for 0, 24, 48, and 72 hours (n = 3). (**I**) Immunoblot analysis of immune response protein activation from samples in H. GAPDH was used as a loading control. (**J**) Percentage of fibroblast cells undergoing *IFNR* TSSG in response to DMSO, 48-hour pIC transfection, 24-hour hydroxy urea (HU) treatment, or 48-hour pIC transfection and 24-hour HU treatment (n = 3). (**K**) Immunoblot analysis of immune response protein activation and the cell cycle factor pRB from samples in J. GAPDH was used as a loading control. (**L**) Percentage of mouse primary T cells undergoing *Ifnr* TSSG in response to 24-hour pIC transfection (n = 3). (**M**) Immunoblot analysis of immune response protein activation from samples in M. GAPDH was used as a loading control. Unless otherwise noted, statistical significance was assessed by a 2-tailed student’s t-test. Black dots represent independent biological replicates; error bars indicate SD.

Here we test the hypothesis that activation of innate immune signaling promotes TSSG of key immune response genes. We report inducible, transient copy number changes at the interferon receptor (*IFNR*) locus on Chr21 (Chr21q22.11). We describe the control of *IFNR* TSSGs by NF-κB in response to exogenous nucleic acids. This signaling cascade culminates in recruitment of the transcription factor RelA to a specific sequence element near the *IFNR* locus that functions as an inducible origin of replication. Removal or suppression of this sequence element blocks generation of *IFNR* TSSGs and targeted recruitment of RelA or epigenetic regulators to the control element is sufficient to induce TSSG production. Genome-wide analysis further identified up to 179 additional loci that share molecular characteristics with *IFNR* locus and may also produce TSSG. Subsequent assessment of six such loci confirmed that recruitment of RelA and DNA replication origin proteins specifies regions that produce TSSGs. Our results describe, for the first time, a signaling cascade and transcription factor-controlled program of inducible origins of replication that produce a transient increase in copy number of key immune-related genes in response to innate immune activation. Our findings reveal an unanticipated regulatory axis of ecDNA production in innate immune activation and provide insights into regulated heterogeneity in innate immune activation.

## RESULTS

### The IFNR locus on chr21 undergoes TSSG in response to innate immune activation

To test the hypothesis that the *IFNR* locus on chr21 undergoes transient site-specific copy number gain (TSSG), we designed two DNA fluorescence *in situ* hybridization (FISH) probes, each spanning ∼200kb: targeting the *IFNR* cluster (red) and a control region ∼2Mb downstream from the cluster (green, **Figure 1B**). We transfected immortalized, but non-transformed diploid fibroblasts for 24 hours with exogenous nucleic acids: the dsRNA mimetic polyinosinic:polycytidylic acid (pIC), as well as ssRNA, dsDNA, and ssDNA. Using DNA FISH, we found a fraction of cells within the population generated an additional copy of the *IFNR* locus, but not the control region, indicating a site-specific, focal copy gain (**Figures 1C** and **1D**, %TSSG is percent of cells with more copies of *IFNR* than control locus, see **Methods** and **Table S1**). Activation of the innate immune response was confirmed via immunoblot for phosphoproteins in the interferon (IFN) and NF-κB signaling cascades as well as the cytoplasmic RNA sensor MDA5 which is common downstream effector of both (**Figures 1E** and **S1A**). Each exogenous nucleic acid increased levels of pTBK1, pRelA, and MDA5, with ssDNA having the most modest effect while still inducing TSSG (**Figures 1D, 1E,** and **S1A**). Similar results were obtained using an immortalized, but non-transformed retinal pigment epithelial (RPE) cell line when transfected with ssRNA or pIC, but not DNA (**Figure 1F**). Immunoblotting confirmed that these RPE cells do not mount an immune response to either dsDNA or ssDNA (**Figures 1G** and **S1B**), likely because they lack expression of cytoplasmic DNA sensors.^56^ When we transfected cells with pIC and assessed TSSG levels by FISH at 24-, 48-, and 72-hour timepoints, we observed that the number of cells with an extra *IFNR* copy was elevated at 24 and 48 hours before returning to baseline levels by 72 hours (**Figure 1H**), indicating these gains are transient in nature and not permanent genetic changes. These changes in *IFNR* copy number were accompanied by activation and resolution of innate immune signaling as measured by pRelA, pIKBα and MDA5 by immunoblot (**Figure 1I**). Furthermore, induction of G1/S cell cycle arrest with hydroxyurea (HU) blocked the formation of *IFNR* TSSG, consistent with a requirement for S phase progression (**Figure 1J** and **K**), a key characteristic of TSSGs.^50-55,57^ Finally, to test whether TSSG of the *IFNR* locus is conserved across species, we isolated primary T cells from mice, activated them with CD3/CD28 costimulation, and transfected them with pIC. When we examined *Ifnr* copy number via FISH, we found elevated copy number in response to pIC-induced innate immune activation (**Figure 1L** and **M**) confirming a conservation of the *IFNR* TSSG in response to innate immune signaling across species.

### NF-κB signaling is required for TSSG of the IFNR locus

To identify potential signaling pathways that regulate TSSG of the *IFNR* locus in an unbiased manner, we performed RNA sequencing on fibroblasts transfected with pIC or dsDNA for 24 hours. Nucleic acid transfection resulted in global changes to the transcriptome with hundreds of differentially expressed genes, exemplified by *TNFAIP3, CCL2,* and *RSAD2* for pIC, and *IFI44L, MX1,* and *TNFRSF10D* for dsDNA (**Figures 2A** and **2B** and **Table S2**). Gene Set Enrichment Analysis (GSEA) of Hallmark Gene Sets identified the IFN Gamma, IFN Alpha, and TNFL Signaling Via NF-κB responses among the most enriched pathways in response to both treatments (**Figure 2C** and **Table S3**). Transcriptome analysis of RPE cells confirmed a robust IFN and NF-κB gene expression response to pIC, but a blunted response to dsDNA, consistent with reduced DNA sensing capability (**Figures S2A-S2C**). We next assessed the relative contributions of each of these innate immune signaling pathways to TSSG formation at the *IFNR* locus using complementary pharmacologic and genetic approaches (**Figure S2D**). To block IFN signaling, we employed the JAK1/2 inhibitor ruxolitinib and found that it had no effect on TSSG induction by pIC,^58-60^ though it did block STAT1 phosphorylation and expression of the ISGs MDA5 and MX1 (**Figures 2D and 2E**). Similar results were obtained for dsDNA (**Figures 2F** and **2G**). Furthermore, using siRNAs to deplete *TBK1* or its effector *IRF3* did not block the induction of TSSG by pIC (**Figures 2H-2K**). In contrast, blockade of NF-κB signaling with the IKKβ inhibitor TPCA1 completely blocked the ability of cells to form TSSG of the *IFNR* locus in response to pIC (**Figures 2L** and **2M**) or dsDNA (**Figures 2N** and **2O**).^61,62^ We confirmed the requirement for NF-κB signaling genetically using siRNAs to deplete *IKK*β and *RelA*, both of which were required for TSSG formation (**Figures 2P-2S**).

**Figure 2.**
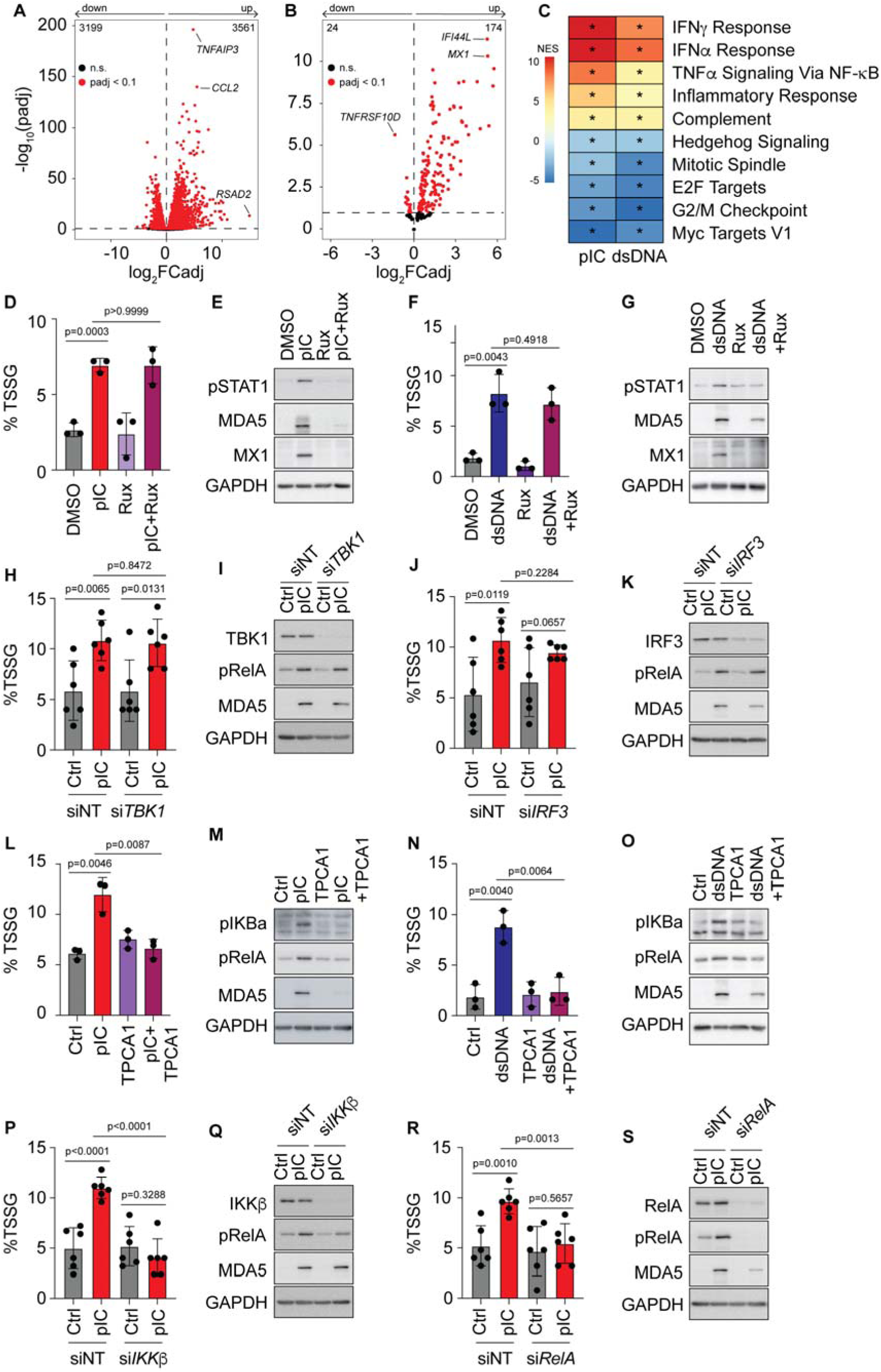
NF-κB is required for TSSG of the *IFNR* locus. (**A**) and (**B**) Volcano plots summarizing transcriptome changes (q<0.1) in fibroblasts following 24-hour transfection of pIC (31.25 ng/mL, **A**) or dsDNA (250 ng/mL, **B**). (**C**) Heatmap of gene-set enrichment analysis (GSEA) of Hallmarks gene sets. Asterisks in the box denote significance (q-value < 0.1) after Benjamini-Hochberg adjustment of *p* values. (**D, F**) Percentage of cells undergoing *IFNR* TSSG in fibroblasts transfected with pIC (D) and dsDNA (F) in the presence or absence of the JAK1/2 inhibitor ruxolitinib (5 μM). Cells were treated with ruxolitinib 1 hour prior to transfection (n =3). (**E, G**) Immunoblot analysis of immune response activation and inhibition for samples in D and F. GAPDH was used as a loading control. (**H, J**) Percentage of cells undergoing *IFNR* TSSG in fibroblasts transfected with non-targeting (NT) or *TBK1* (H) and *IRF3* (J)-targeting siRNA for 48 hours prior to pIC transfection. Results represent 3 independent transfections of 2 different siRNAs. (**I, K**) Immunoblot analysis of siRNA target knockdowns and immune response activation from H, and J. GAPDH was used as a loading control. (**L, N**). Percentage of cells undergoing *IFNR* TSSG in fibroblasts transfected with pIC (H) and dsDNA (J) in the presence or absence of the IKKβ inhibitor TPCA1 (2 μM). Cells were treated with TPCA1 1 hour prior to transfection (n =3). (**M, O**) Immunoblot analysis of immune response activation and inhibition for samples in L and N. GAPDH was used as a loading control. (**P, R**) Percentage of cells undergoing *IFNR* TSSG in fibroblasts transfected with non-targeting (NT) or IKKβ (*IKBKB*) (P), and *RELA* (R)-targeting siRNA for 48 hours prior to pIC transfection. Results represent 3 independent transfections of 2 different siRNA. (**Q, S**) Immunoblot analysis of siRNA target knockdowns and immune response activation from P, and R. GAPDH was used as a loading control. Statistical significance was assessed by a 2-tailed student’s t-test. Black dots represent independent biological replicates. Error bars indicate SD.

### TNF**α** is sufficient to induce TSSG at the IFNR locus

Based on our observations that IKKβ and RelA are required for TSSG at the *IFNR* locus, we next asked whether TNFα, a canonical activator of the NF-κB pathway, is sufficient to drive this process. TNFα treatment induced TSSG at the *IFNR* locus within 24 hours that persisted for 48 hours, but were resolved by 72 hours, consistent with TSSG of the *IFNR* locus (**Figures 3A** and **3B**). Notably, TNFα also induced *Ifnr* TSSG in primary mouse T cells (**Figures 3C** and **3D**) consistent with conservation of TSSG of the *IFNR* locus in response to TNFL across species. Next, we assessed the mechanism of TNFα induction of *IFNR* TSSG using genetics. We found that knockdown of *TBK*1 and *IRF3* did not block *IFNR* TSSG (**Figures 3E**-**3H**). As was the case for pIC, we observed that knockdown of either *IKK*β or *RelA* blocked the indication of TSSG by TNFα, indicating that NF-κB signaling is both necessary and sufficient for this process (**Figures 3I**-**3L**). Consistent with this, inhibition of IKKβ by TPCA1 also blocked *IFNR* TSSG formation (**Figures 3M and 3N**).

**Figure 3.**
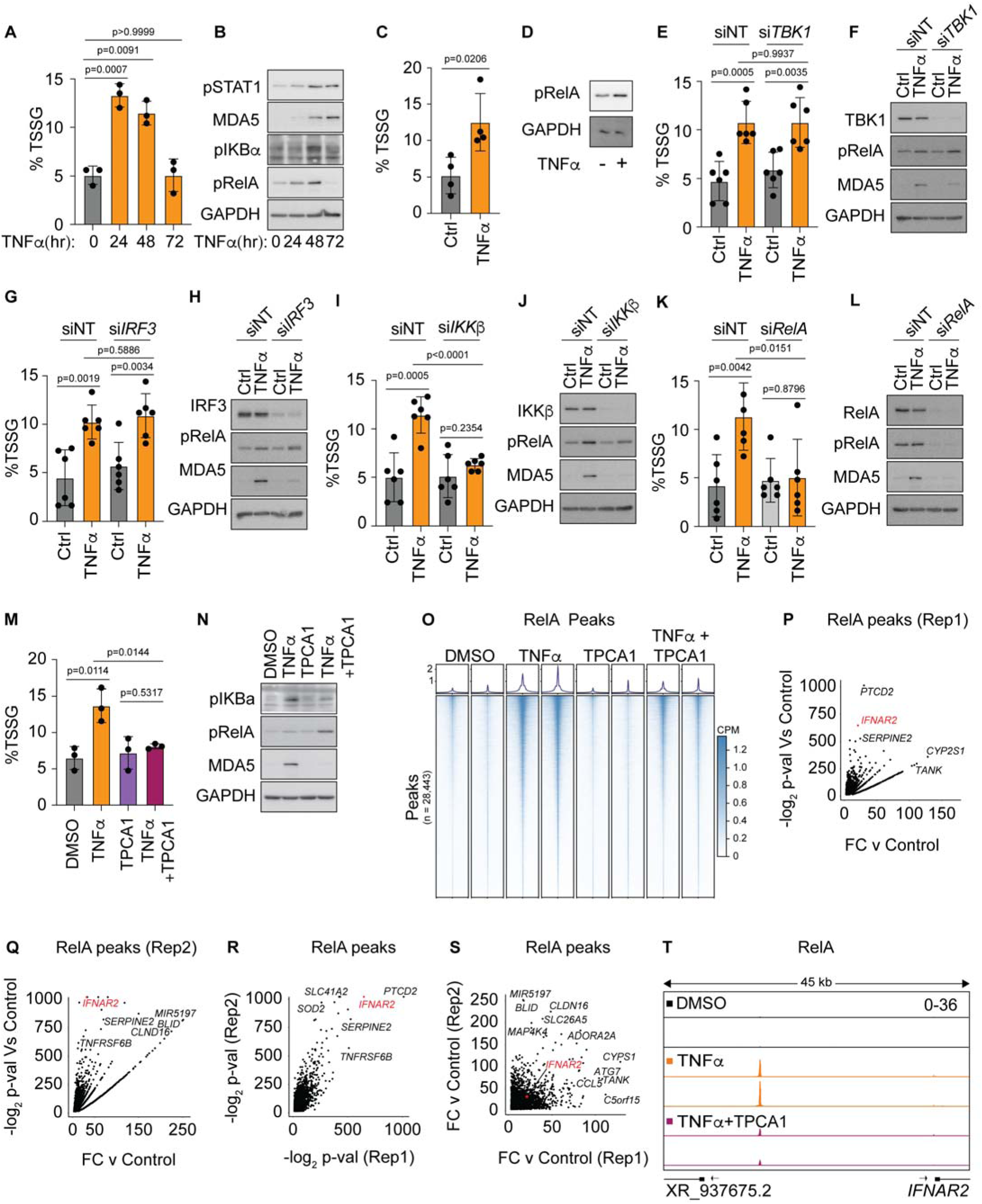
TNF**α** drives TSSG of *IFNR* locus. (**A**) Percentage of fibroblasts undergoing *IFNR* TSSG treated with TNFα (5 ng/mL) for 0, 24, 48, and 72 hours (n = 3). (**B**) Immunoblot analysis of immune response protein activation from samples in A. GAPDH was used as a loading control. (**C**) Percentage of mouse primary T cells undergoing *Ifnr* TSSG in response to 24-hour TNFα treatment (n = 4). (**D**) Immunoblot analysis of immune response protein activation from samples in C. GAPDH was used as a loading control. (**E, G, I, K**) Percentage of cells undergoing *IFNR* TSSG in fibroblasts transfected with non-targeting (NT) or *TBK1* (E), *IRF3* (G), IKKβ (*IKBKB*) (I), and *RELA* (K)-targeting siRNA for 48 hours prior to TNFα treatment. Results represent 3 independent transfections of 2 different siRNAs. (**F, H, J, L**) Immunoblot analysis of siRNA target knockdowns and immune response protein activation from E, G, I, and K. GAPDH was used as a loading control. (**M**) Percentage of fibroblasts undergoing *IFNR* TSSG for biological replicates utilized for all TNFα treated CUT&RUN experiments (n = 3). Two replicates were used for CUT&RUN. (**N**) Immunoblot analysis of pIKBα and RelA for CUT&RUN samples in **M.** (**O**) Heatmaps surrounding HOMER peak calls from two biological RelA CUT&RUN experiments for the indicated treatment. Each row represents a peak that was called in at least one treatment condition against a matched IgG control. Overlapping peaks across treatment conditions were merged. Columns range from -2.5 kb to 2.5 kb from peak center with 50 bp bins. (**P-Q**) Fold change vs -log_2_ p-value plot for inducible RelA HOMER peak calls in TNFα treated cells for each individual replicate. (**R**) Scatter plot representing -log_2_ p-values from both biological replicates of TNFα RelA CUT&RUN experiments. (**S**) Scatter plot of fold changes from both biological replicates of TNFα RelA CUT&RUN experiments. (**T**) RelA binding site upstream of *IFNAR2*. CUT&RUN tracks from two biological replicates treated as indicated. Scale in upper right applies to all tracks and is CPM normalized. Unless otherwise noted, statistical significance was assessed by a 2-tailed student’s t-test. Black dots represent independent biological replicates. Error bars indicate SD.

To further explore the mechanism of NF-κB-dependent *IFNR* TSSG, we next performed Cleavage Under Target & Release Using Nuclease (CUT&RUN), to examine the genome-wide binding of RelA in response to TNFα treatment with or without TPCA1. We observed increased RelA recruitment genome wide following TNFα treatment (**Figure 3O**). Notably, we identified a significant, reproducible RelA binding site upstream of *IFNAR2* that was strongly inducible by TNFα and blocked by TPCA1 (**Figures 3P-3T**). In parallel, we performed CUT&RUN on pIC treated fibroblasts and observed (**Figures S3A-S3C**) with strong reproducible recruitment of RelA that was ablated by TPCA1 treatment (**Figures S3D-S3H**). We identified binding of RelA to the same element for cells treated with pIC (**Figures S3D-S3H**). These findings are consistent with NF-κB signaling driving a transient copy number increase of the *IFNR* locus in response to innate immune signaling.

### NF-κB signaling drives epigenetic changes at the IFNR locus

Previously identified TSSGs are controlled epigenetically through changes in the local chromatin environment.^50-55,63^ To identify potential regulators of the *IFNR* TSSG, we investigated changes in chromatin modifications in response to NF-κB signaling by TNFα treatment. We profiled histone modifications associated with TSSGs, ecDNA and DNA replication including H3K4me1, H3K4me3, H3K9me1, H3K9me3, H3K27me3, H3K36me3, H3K37me1, H4K20me1, and H3K27ac.^47,48,50-55,64-66^ If epigenetic changes were associated with TSSG production we expected treatment with TNFα to increase or decrease signal at the IFNR locus and particularly at the RelA binding site. We did not observe any dynamic regulation of H3K9me3, H3K9me1, H3K36me3, H3K37me1, H3K27me3, and H4K20me1 at the RelA binding site upstream of *IFNAR2* (**Figure 4**), nor were these modifications enriched at the locus (**Figure 4A, C, E, G, I, and K**), though all were detected globally (**Figures 4B, D, F, H, J, and L**). Notably, the *IFNAR2* RelA binding site exhibited reduced levels of H3K27me3 compared to surrounding chromatin, but this was not regulated by TNFα (**Figures 4I and 4J**). In contrast, we did observe a TNFα- and NF-κB-dependent increase in H3K4me1 and H3K27ac at the *IFNAR2* RelA binding site (**Figures 4M**-**4P**). H3K27ac is primarily mediated by the highly similar acetyltransferases p300 and CBP.^67^ Consistent with a role for p300/CBP and H3K27ac in regulating *IFNR* TSSG, pharmacological inhibition of the chromatin binding of p300/CBP using the p300/CBP bromodomain inhibitor SGC-CBP30 (2 µM) completely blocked TSSG formation (**Figures 4Q** and **4R**).^68,69^ These results suggest the RelA binding site and associated epigenetic changes contribute to regulation of the *IFNR* TSSG.

**Figure 4.**
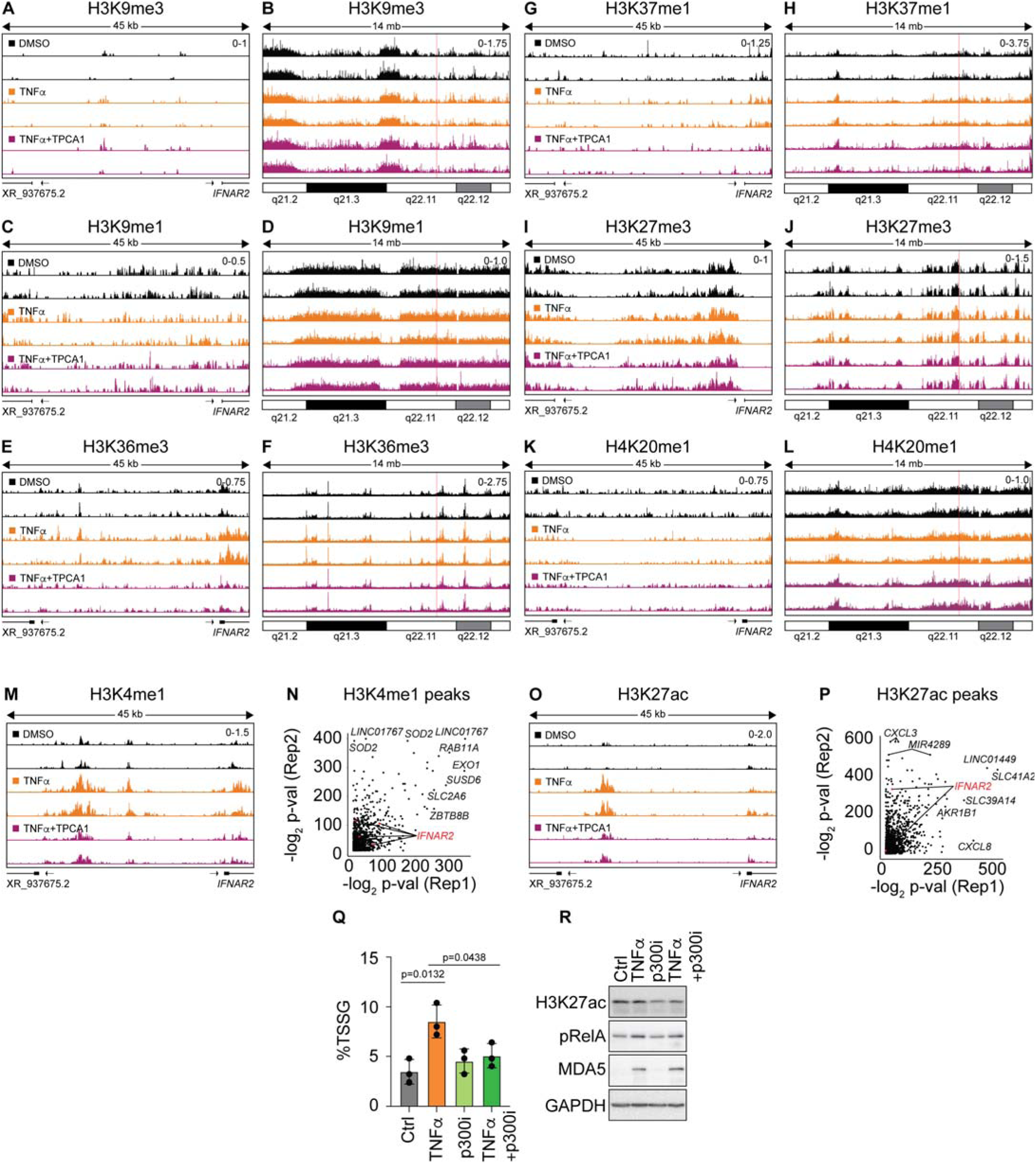
Epigenetic Regulation of an *IFNR* TSSG. (**A** to **L**) CUT&RUN tracks for two biological replicates treated with TNFα (5 ng/mL) for 24 hours for H3K9me3, H3K37me1, H3K9me1, H3K36me3, H4K20me1 and H3K27me3. For each mark, a zoomed 45kb view of the RelA binding site and a 14Mb view are provided. Vertical red line indicates location of the 45kb region. (**M**) H3K4me1 CUT&RUN tracks from two biological replicates of fibroblasts treated with TNFα (5 ng/mL) for 24 hours, with or without TPCA1 pretreatment. (**N**) Scatter plot representing -log_2_ p-values of H3K4me1 peaks from both biological replicates of CUT&RUN following TNFα treatment. (**O**) H3K27ac CUT&RUN tracks from two biological replicates of fibroblasts treated with TNFα (5 ng/mL) for 24 hours, with or without TPCA1 pretreatment. (**P**) Scatter plot representing -log_2_ p-values of H3K27ac peaks from both biological replicates of CUT&RUN following TNFα treatment. (**Q**) Percentage of fibroblasts undergoing *IFNR* TSSG from treated for 24 hours with TNFα in the presence or absence of the p300/CBP inhibitor SGC-CBP30 (2 μM). Cells were treated with the inhibitor 1 hour prior to TNFα treatment (n = 3). (**R**) Immunoblot analysis of immune response activation and inhibition for samples in Q. GAPDH was used as a loading control. Unless otherwise noted, statistical significance was assessed by a 2-tailed student’s t-test. Black dots represent independent biological replicates. Error bars indicate SD.

### The RelA binding site upstream from IFNAR2 is a TSSG Control Element

Next, we asked whether the inducible RelA binding site upstream of *IFNAR2* is required for the formation of the *IFNR* TSSG. Using CRISPR-RNP editing with two independent guide RNAs, we targeted a 259 bp segment of DNA centered on the inducible *IFNAR2* RelA binding site (**Figure S4A**). Notably, we were unable to obtain homozygous deletions for this intergenic region, however, we were able to generate and isolate numerous clones harboring heterozygous deletions. Three heterozygous clones were pooled (Het-RNP) and compared to a pool of three clones that received the RNP-Cas9 transfection but retained both wild-type alleles at the *IFNR*-RelA peak location (WT-RNP) (**Figure S4B**). TSSG induction at the *IFNR* locus was significantly decreased in the Het-RNP pool (**Figure 5A**). As expected, the Het-RNP pool showed a reduction in the interaction of RelA at the binding site upstream of *IFNAR2* following TNFα treatment (**Figures 5B-D**). These data indicate that the RelA binding site upstream of *IFNAR2* is required for *IFNR* TSSG and thus functions as a TSSG Control Element (TCE). Consistent with a role in TSSG production, the Het-RNP pool not only exhibited reduced binding of RelA but was accompanied by decreased levels of H3K4me1 and H3K27ac (**Figures 5E-J**). To further establish the importance of the RelA binding site in controlling TSSG of the *IFNR* locus, we attempted to suppress the control element through recruitment of a dCas9-EZH2 fusion.^70^ We designed six guide RNAs targeting the 400 bp flanking the *IFNR* RelA binding site (**Figure 5K**). Recruitment of dCas9-EZH2 to the *IFNR* TCE, but not a control genomic locus, was sufficient to ablate TSSG production at the *IFNR* locus following TNFα treatment (**Figure 5L**). The ability of dCas9-EZH2 to suppress the *IFNR* TSSG through either steric occlusion of the RelA binding site or EZH2-induced heterochromatinization further underscores the importance of the RelA binding site. Taken together our analysis supports a role for the RelA binding site as a TCE.

**Figure 5.**
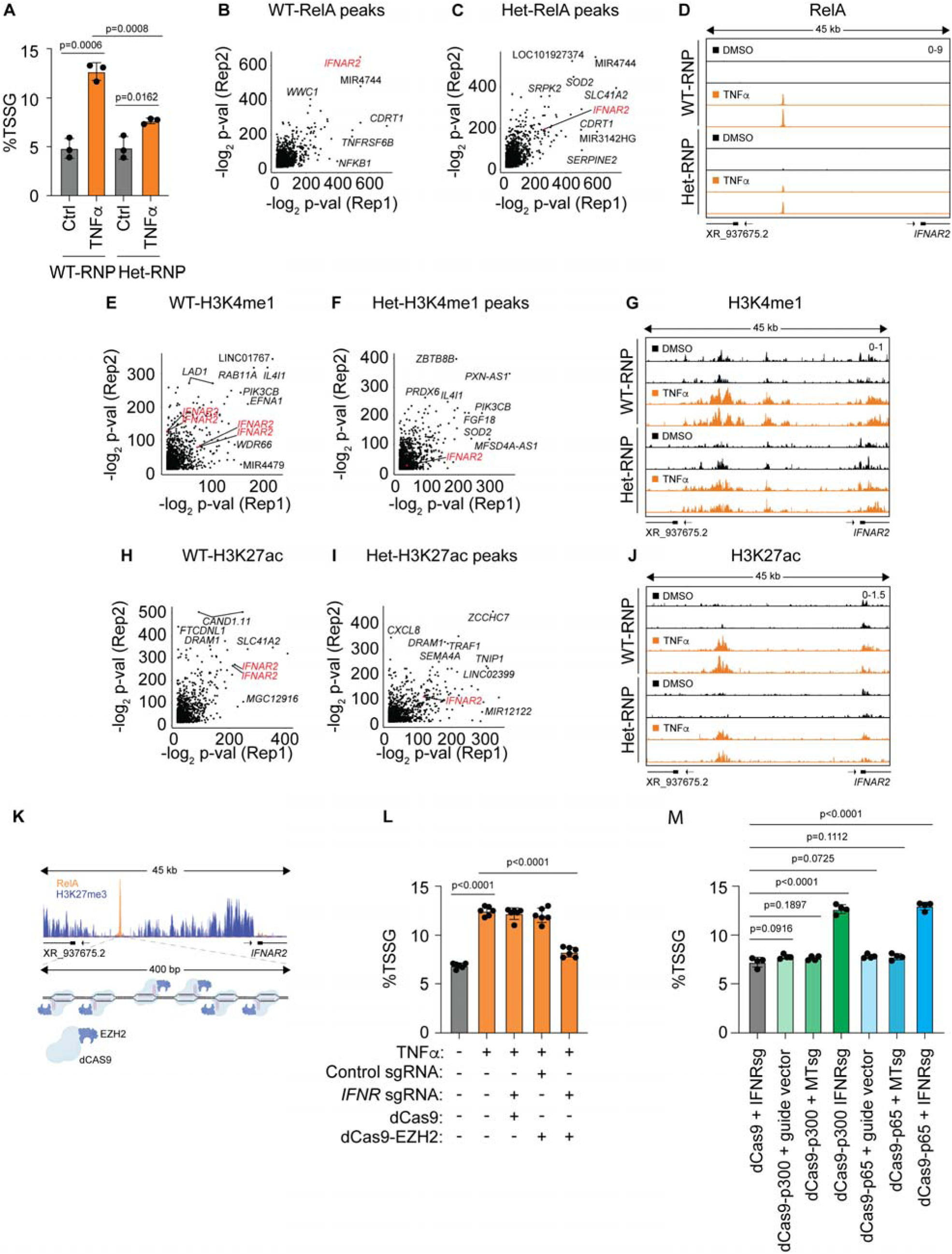
The RelA binding site functions as a TSSG Control Element. (**A**) Percentage of fibroblasts undergoing *IFNR* TSSG in WT-RNP and Het-RNP fibroblasts treated with 5 ng/mL TNFα for 24 hours (n = 3). (**B**) Scatter plot representing -log_2_ p-values from RelA CUT&RUN peaks from both biological replicates of WT-RNP TNFα treatment. (**C**) Scatter plot representing -log_2_ p-values from RelA CUT&RUN peaks from both biological replicates of Het-RNP TNFα treatment (**D**) RelA binding at the TCE. CUT&RUN tracks from two biological replicates in WT-RNP and Het-RNP fibroblasts treated as indicated. (**E**) Scatter plot representing -log_2_ p-values from peaks from both biological replicates of WT-RNP TNFα treated H3K4me1 CUT&RUN. (**F**) Scatter plot representing -log_2_ p-values from peaks from both biological replicates of Het-RNP TNFα treated H3K4me1 CUT&RUN. (**G**) H3K4me1 at the TCE. H3K4me1 CUT&RUN tracks from two biological replicates of WT-RNP and Het-RNP fibroblasts treated as indicated. (**H**) Scatter plot representing -log_2_ p-values from peaks from both biological replicates of WT-RNP TNFα treated H3K27ac CUT&RUN. (**I**) Scatter plot representing -log_2_ p-values from peaks from both biological replicates of Het-RNP TNFα treated H3K27ac CUT&RUN. (**J**) H3K27ac at the TCE. H3K27ac CUT&RUN tracks from two biological replicates of WT-RNP and Het-RNP fibroblasts treated as indicated. (**K**) Schematic of dCas9-EZH2 targeting experiment. Six independent guide RNAs targeted dCas9 alone or dCas9-EZH2 to the TCE or another genomic locus (control sgRNA). (**L**) Percentage of cells undergoing *IFNR* TSSG in fibroblasts transfected with the indicated constructs and treated with 5 ng/mL TNFα. Results represent 6 independent transfections of sgRNA. (**M**) Percentage of cells undergoing *IFNR* TSSG in fibroblasts transfected with the indicated constructs. Results represent 6 independent transfections Unless otherwise noted, statistical significance was assessed by a 2-tailed student’s t-test. Black dots on bar graphs represent independent biological replicates. Error bars indicate S.D.

We next asked if the TCE could also function to induce TSSG of the *IFNR* locus in the absence of global changes caused by innate immune activation. Using the same guide RNAs used to suppress the locus with dCas9-EZH2, we tested whether recruitment of RelA or p300 was sufficient to induce *IFNR* TSSG. We used dCas9 fusion constructs containing the RelA activation domain or the p300 histone acetyltransferase (HAT) domain.^71,72^ Recruitment of either RelA or the p300 histone acetyltransferase domain to the TCE, but not a control genomic locus, was sufficient induce *IFNR* TSSG (**Figure 5M**). Together our results demonstrate the importance of the TCE, recruitment of RelA and epigenetic control through p300 in regulating TSSG of the *IFNR* locus.

### NF-κB activation drives a program of inducible DNA replication origins characterized by MCM2 binding

To determine how the TCE regulates *IFNR* TSSG, we asked if this element could serve as an inducible origin of replication. We performed CUT&RUN for the replication initiation factor minichromosome maintenance 2 (MCM2) whose recruitment is indicative of a licensed origin of replication.^48^ Interestingly, TNFα had little to no effect on global MCM2 binding sites (**Figure 6A** and **Table S4**). However, we identified 1,359 MCM2 peaks induced upon TNFα treatment that were consistent across replicates (**Figures 6B-E** and **S5A** and **Table S5)**. Analysis of the genes near these loci using GSEA revealed strong enrichment for the Hallmark Interferon Gamma and Interferon Alpha gene sets (**Figure 6F and Table S6**). TNFα treatment induced a strong MCM2 peak coincident with the TCE upstream of *IFNAR2* (**Figure 6G**). This inducible MCM2 peak is partially blocked by pretreatment with TPCA1, suggesting a requirement for IKKβ signaling and RelA association prior to MCM2 recruitment (**Figure 6G**). Furthermore, heterozygous knockout of the TCE upstream of *IFNAR2* (Het-RNP) reduced the levels of MCM2 recruitment to this locus upon TNFα treatment (**Figure 6H**).

**Figure 6.**
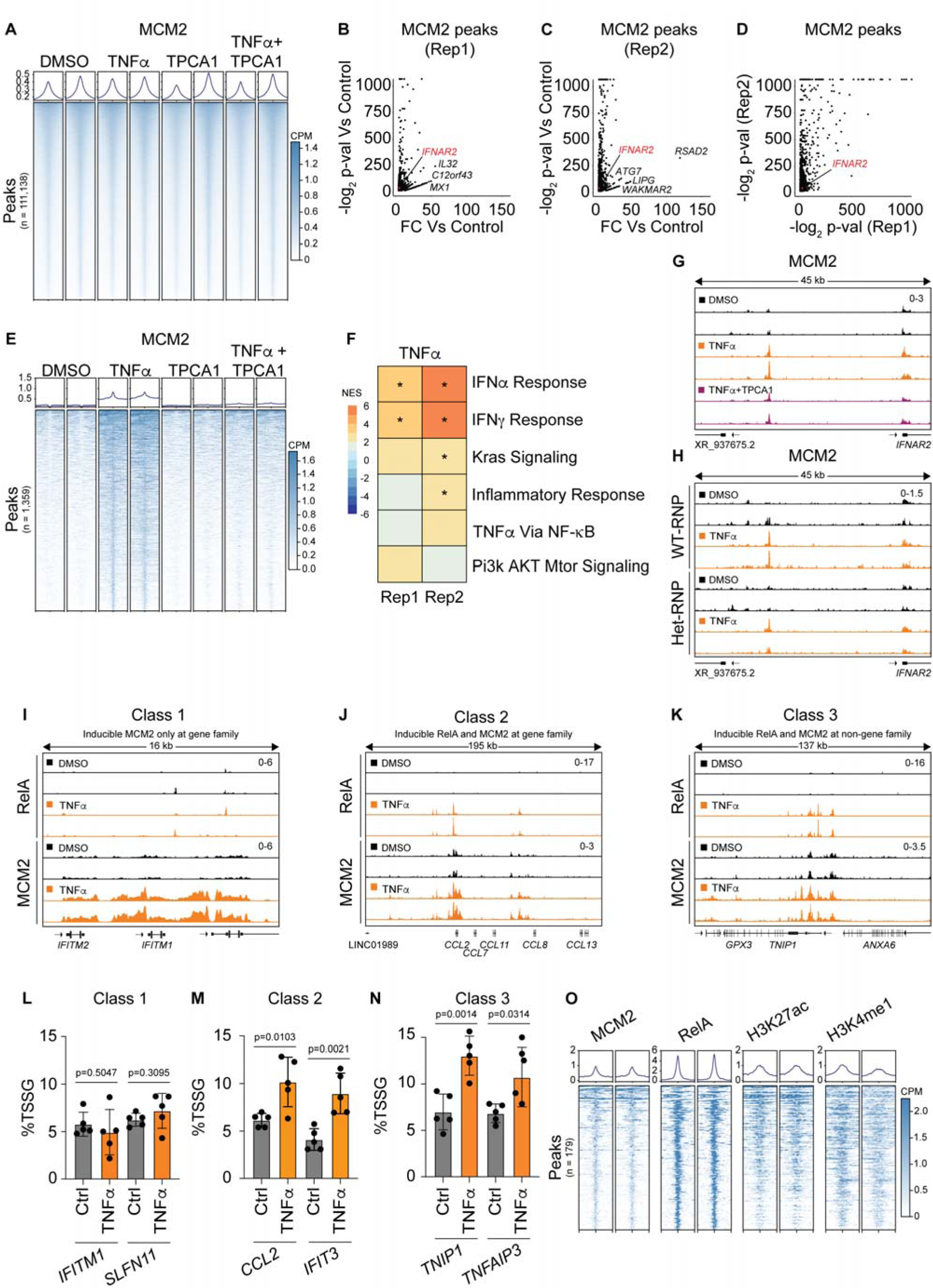
Innate immune activation drives a program of TSSGs at immune response genes. (**A**) Heatmaps of MCM2 signal surrounding HOMER peak calls from two biological replicates of MCM2 CUT&RUN for the indicated treatment. Each row represents a peak that was called in at least one treatment condition against a matched IgG control. Overlapping peaks across treatment conditions were merged. Columns range from -2.5 kb to 2.5 kb from peak center with 50 bp bins. (**B** and **C**) Fold change vs -log_2_ p-value plot for MCM2 peaks called by HOMER in TNFα treated cells for each independent replicate. (**D**) Scatter plot representing -log_2_ p-values from peaks from both biological replicates of TNFα treated MCM2 CUT&RUN peaks called by HOMER from B and C. (**E**) Heatmaps of MCM2 signal surrounding HOMER peak calls from TNFα vs DMSO treatments from two biological replicates of MCM2 CUT&RUN for the indicated treatment. Each row represents a peak that was called in both replicates for TNFα against its DMSO associated control. Columns range from -2.5 kb to 2.5 kb from peak center with 50 bp bins. (**F**) Heatmap of gene-set enrichment analysis (GSEA) of the nearest gene to TNFα-inducible MCM2 peaks using Hallmark gene sets. Asterisks in the box denote significance (q-value < 0.1) after Benjamini-Hochberg adjustment of *p* values. (**G**) MCM2 at the TCE. MCM2 CUT&RUN tracks of the TCE from two biological replicates treated as indicated. (**H**) MCM2 at the TCE. MCM2 CUT&RUN tracks of the TCE from two biological replicates in WT-RNP and Het-RNP fibroblasts treated as indicated. (**I**) MCM2 and RelA CUT&RUN tracks from two biological replicates of a Class-1 site (inducible MCM2 only at repetitive gene family (e.g., *IFITM1* locus)) treated as indicated. (**J**) MCM2 and RelA CUT&RUN tracks from two biological replicates of a Class-2 site (inducible RelA and MCM2 at repetitive gene family (e.g., *CCL2* locus)) treated as indicated. (**K**) MCM2 and RelA CUT&RUN tracks from two biological replicates of a Class-3 site (inducible RelA and MCM2 at a non-repetitive gene family locus (e.g. *TNIP1*)) treated as indicated. (**L-N**) Percentage of fibroblasts undergoing TSSG treated with 5 ng/mL TNFα for 24 hours for class I (I, n = 5), class II (**J, n= 5**), and class III (**K, n = 5**) genes. (**O**) Heatmaps from two replicates of MCM2, RelA, H3K27ac, and H3K4me1 CUT&RUN after TNFα treatment for 179 loci with TNFα-inducible overlapping MCM2 and RelA binding. Note only the two replicates of TNFα treatment are shown. Columns range from -2.5kb to 2.5kb from peak center with 50 bp bins. Unless otherwise noted, statistical significance was assessed by a 2-tailed student’s t-test. Black dots on bar graphs represent independent biological replicates. Error bars indicate S.D.

We reasoned that RelA binding and inducible MCM2 recruitment may define other loci that also produce TSSG following TNFα stimulation. Additionally, we considered whether repetitive gene family loci, may be more likely to generate TSSG since the *IFNR* locus, *MT* locus, and *HSATII* locus, all generate TSSGs.^51,55^ We therefore classified inducible MCM2 peaks (n=1,359) into three classes of genomic regions: class 1 - inducible MCM2-only (n=1,180); class 2 - inducible MCM2 and RelA at repetitive gene families (n=18); and class 3 - inducible MCM2 and RelA at non-repetitive gene families (n=161) (**Figures 6I-6K** and **Table S7**). We developed DNA FISH probes to representative loci from each class and prioritized testing potential TSSG sites at genes related to interferon and TNFα/NF-κB signaling. At the Class 1 gene *IFITM1*, we observed strong MCM2 recruitment in the absence of RelA binding, but this locus failed to produce TSSG. A similar result was observed at *SLFN11*, another Class 1 locus that lacked inducible RelA binding (**Figures 6I** and **6L**). However, the Class 2 locus surrounding *CCL2* exhibited recruitment of both RelA and MCM2 and produced TSSGs in response to TNFα treatment, as did the Class 2 loci at *IFIT3* and *IL1A* (**Figures 6J** and **6M** and **S5B**). In addition, we also observed coincident RelA and MCM2 recruitment and TSSG at the Class 3 loci near *TNIP1* and *TNFAIP3* (**Figures 6K** and **6N**) suggesting that both RelA and MCM2 recruitment, but not a repetitive gene family locus is required for innate immune-driven TSSGs. Finally, we assessed the chromatin landscape of the 179 Class 2 and Class 3 peaks with concurrent MCM2 and RelA recruitment near 165 genes in the innate immune TSSG program (**Table S7**). We found that the majority of these peaks were marked by H3K27ac and H3K4me1, including the Class 2 locus *IFIT3* and the Class 3 locus *TNIP1*, as was the case for the *IFNR* locus (**Figure 6O** and **S5C-D**). Our results describe a program of TSSG production driven by innate immune signaling requiring NF-κB.

## DISCUSSION

Here we describe a program of inducible DNA replication origins specified by NF-κB signaling that produce transient focal gene amplifications in response to innate immune signaling. The program is regulated by RelA and epigenetic changes in close proximity to inducible origins of replication marked by MCM2. We provide the first evidence for a coordinated response to cell signaling that generates additional ecDNA copies of multiple genomic regions important in the response to that signal.

In higher metazoans it has become apparent that origins of replication are frequently associated with promoters of active genes and are thus also enriched in transcription factor binding sites.^48,73,74^ Previous reports have established a role for transcription regulatory regions in marking origins of replication, such as the requirement for the locus control region (LCR) of the β-globin locus for firing of the distal β-globin replication origin.^75^ Surprisingly, we find that transcription factors, in this case RelA, can direct targeted gene amplification through inducible origin licensing in response to activation of a signaling pathway. This completely unexpected finding reveals a role for sequence specific DNA binding proteins in the control of gene amplification in metazoans. Whether and how transcription factors might facilitate inducible origins remains unclear. It is possible RelA directly interacts with components of the replication machinery to facilitate their recruitment or promotes establishment of a permissive chromatin environment at inducible replication origins. Consistent with this idea, we have established that targeted recruitment of RelA or the HAT p300 can generate TSSGs in the absence of innate immune signaling. Recent reports further support a global role for H3K4me1 in regulating origin firing particularly in late replicating regions.^76^

The program of reversible gene amplifications controlled by innate immune signaling we describe here raises the interesting question of how this might contribute to cellular phenotypic heterogeneity. The production of many cytokines and their receptors including interferons occurs in a small fraction of cells in a population and in response to innate immune signaling.^14-19^ Single cell RNA sequencing has further revealed that fewer than 25% of the cells in fibroblasts or mononuclear cells induce transcription of cytokines.^15^ This is consistent with the rate of 10 to 15% off cells producing TSSGs of any given loci in our studies. It is possible that TSSG production contributes to the cellular genotypic and phenotypic heterogeneity in response to innate immune signaling. Thus, even a seemingly low fraction of cells producing transient gene amplifications may exhibit strong biological impact across millions of dividing cells experiencing an innate immune response. Furthermore, TSSGs of the epidermal growth factor receptor (EGFR), at one extra copy in ∼15% of cells, renders cells more responsive to EGF treatment and resistant to the EGFR inhibitor gefitinib supporting a role for low copy and heterogenous effects of ecDNA on cellular responses.^53^ It will be interesting to determine if a cell produces TSSG from all the immune loci or if different cells in the population have a different complement of innate immune-related TSSGs. This could provide an array of phenotypic heterogeneity in tissue response to pathogens. Furthermore, understanding how programs of TSSG production are regulated is likely to provide insights into heterogeneity in drug response and genomic instability in cancer and raises the possibility that the heterogeneity is a regulated and conserved biologically relevant process.

We have described here the first sequence specific control element for TSSGs, a TCE, that is required for TSSG production in response to innate immune signaling. Identification of this element provides a means to control production of specific TSSG events without disrupting the global response to environmental signaling, using genetic manipulation or Cas9-tethering experiments, for example. Consistent with this idea, targeted recruitment of the activation domain of RelA or the HAT domain of p300 was sufficient to induce TSSG at the IFNR locus (**Figure 5M**). This will allow the future assessment of the role of TSSGs in biological stress responses. Further, the ability to identify TCEs through the inducible recruitment of activated transcription factors colocalizing with MCM2 in response to stress should enable more rapid identification of programs of ecDNA produced in other developmental, stress and disease contexts. It is likely that the use of inducible origins in response to stress is not restricted to the NF-κB pathway as with previously identified TSSGs produced in response to hypoxia and metal stress.^50,55^

Previous work with TSSGs has suggested that under the right conditions they can become integrated into chromosomes and selected for as permanent genetic changes, particularly in the presence of prolonged exposure and chronic signaling.^55,63^ The presence of 18 loci with gene families in our analyses suggests the possibility that TSSG production during conditions of prolonged stress could contribute to targeted gene duplication events and development of repetitive gene families. It would be interesting to ascertain if continual, prolonged innate immune signaling leads to amplification and evolution of repetitive gene families. It should be noted that increasing the copy number of the *IFNR* locus, as is the case for Trisomy 21, has recently been linked to the interferonopathy of Down Syndrome and a number of co-occurring conditions which affect that population.^60,77-82^

Taken together, our findings highlight a completely unanticipated program of TSSG and ecDNA production in response to a biological signaling pathway with the potential for wide-ranging roles in stress response and developmental programs.

## RESOURCE AVAILABILITY

### Lead contact

Further information and requests for resources and reagents should be directed to and will be fulfilled by the lead contact, Kelly Sullivan.

### Materials availability

This study did not generate any unique materials.

## ACKNOWLEDGMENTS

This work was supported primarily by NIH grants R01AI145988 (JCB and KDS). Additional funding was provided by NIH grants P30CA046934, R35GM128720 (JCB), the Linda Crnic Institute for Down Syndrome, the Global Down Syndrome Foundation, the Anna and John J. Sie Foundation, the University of Colorado School of Medicine, and the Boettcher Foundation.

## AUTHOR CONTRIBUTIONS

ML – Investigation, Visualization, Methodology, Writing; JW– Investigation, Visualization, Methodology, Writing; MDG – Formal Analysis, Visualization, Data Curation, Writing – review & editing; NB – Resources, Investigation, LD – Investigation; JCB – Conceptualization, Supervision, Funding Acquisition, Visualization, Writing; KDS – Conceptualization, Supervision, Funding Acquisition, Visualization, Writing.

## DECLARATION OF GENERATIVE AI AND AI-ASSISTED TECHNOLOGIES

No generative AI was used in the preparation of this manuscript.

## SUPPLEMENTAL INFORMATION

Materials and Methods

Figs. S1 to S5

Tables S1-S12

## KEY RESOURCES TABLE

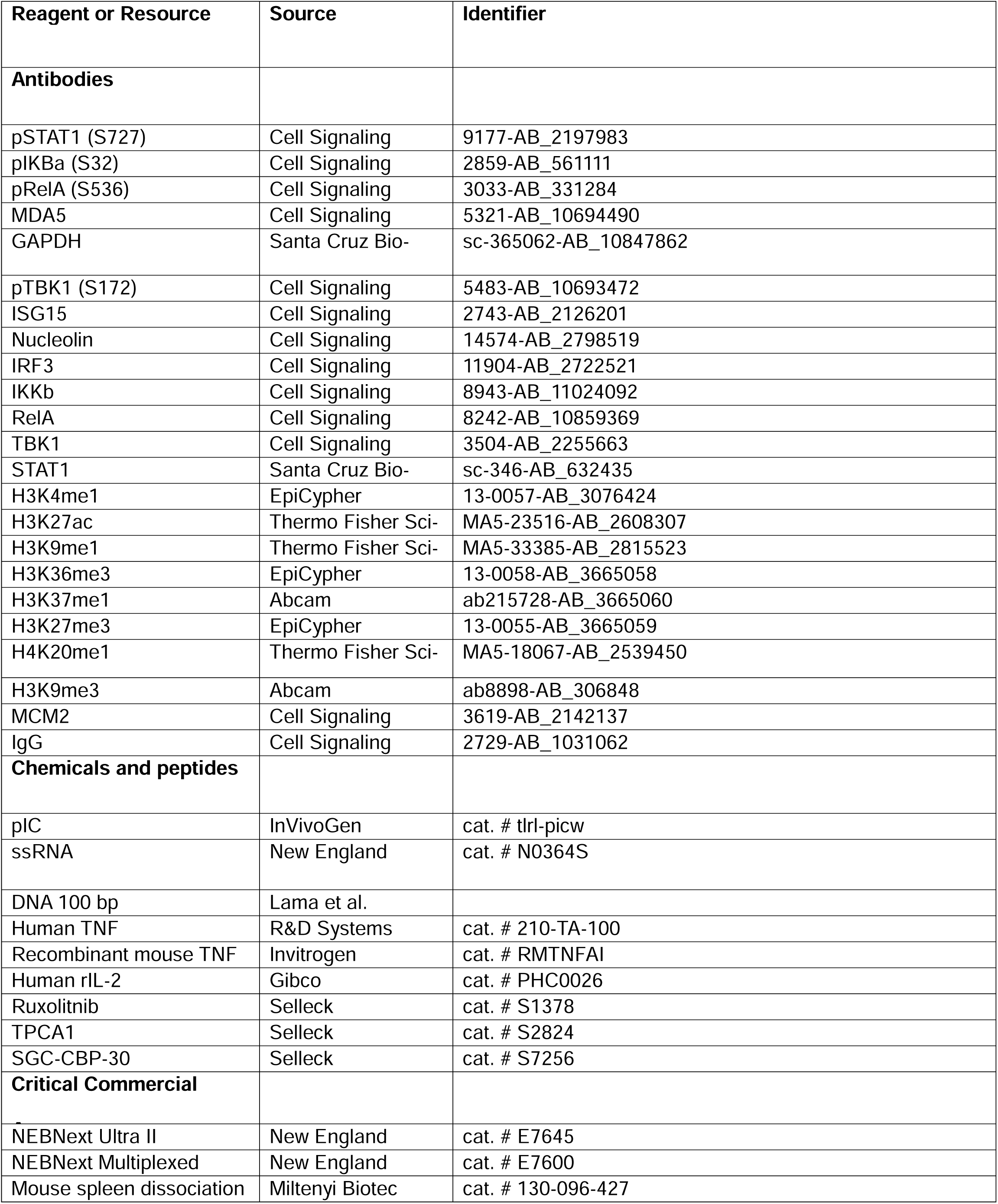
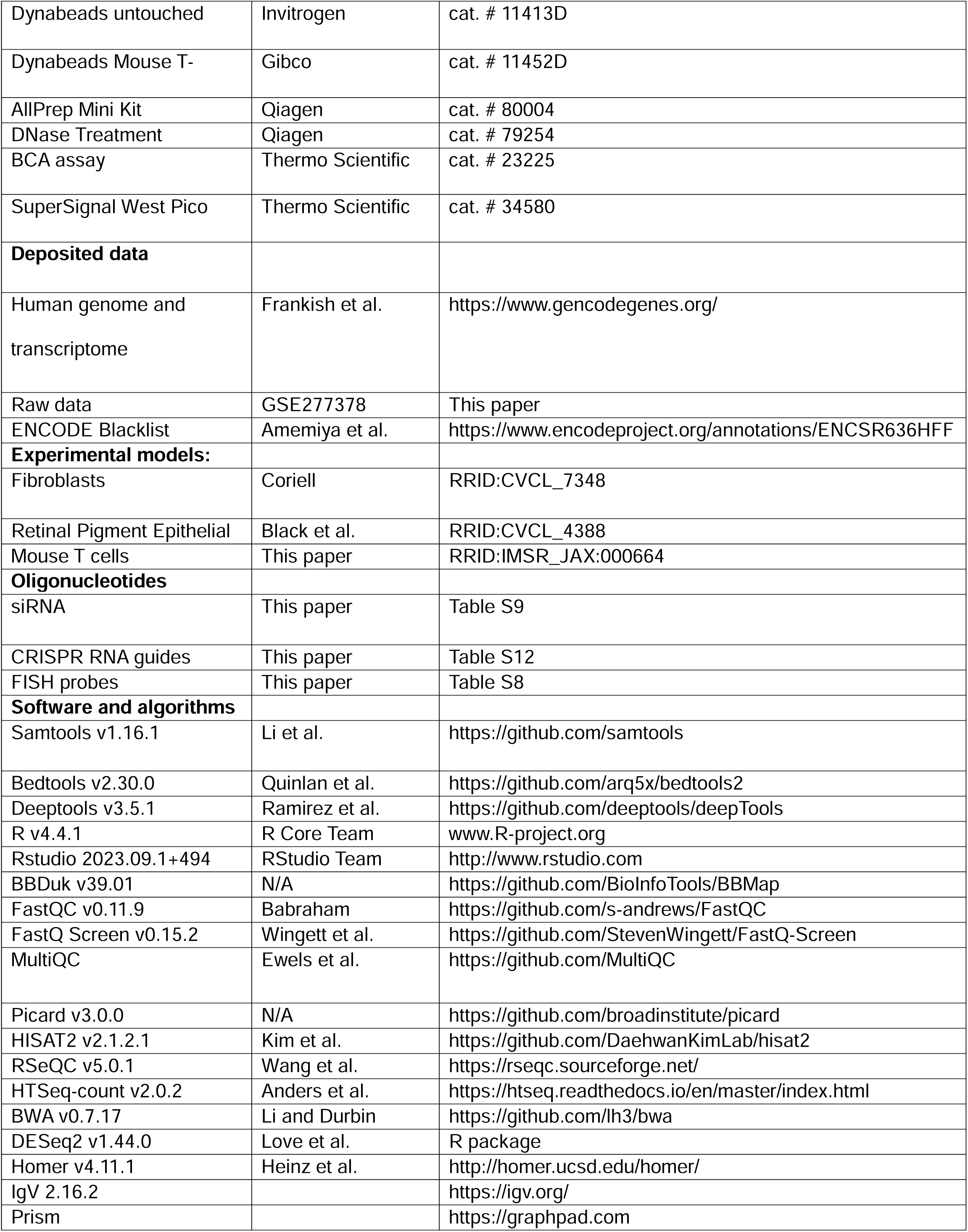
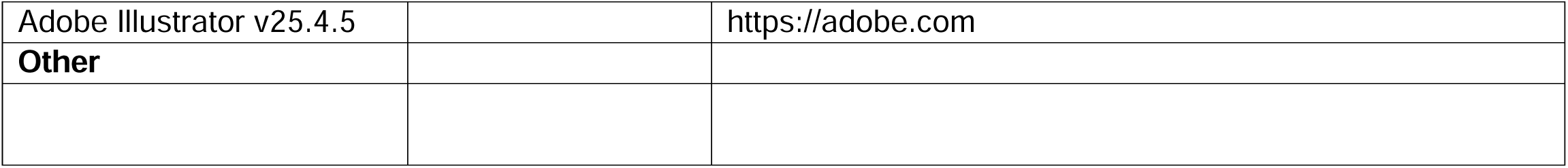

## EXPERIMENTAL MODEL AND STUDY PARTICIPANT DETAILS

### Human Cell culture

Fibroblasts were acquired from Coriell (GM02036, Female Skin Fibroblast) and cultured in DMEM (Thermo Fisher Scientific, cat# 11995-065) supplemented with 10% fetal bovine serum (Peak Serum, cat # PS-FB3) and antibiotic-antimycotic (Anti-Anti, Thermo Fisher Scientific, cat #15240062). RPE cells were obtained from Dr. Johnathan Whetstine and cultured in DMEM (Thermo Fisher Scientific, cat #11995-065) supplemented with 10% FB Essence (VWR, cat# 10803-034), 1x Pen/Strep (Corning, cat# 30-002-Cl) and 2 mM L-Glutamine (Gibco, cat# 25030-081). All cells were cultured in humidified incubators with 5% CO_2_. Cell lines were routinely observed to be mycoplasma free during DNA FISH experiments.

### Mouse T Cell Culture

Experiments were approved by the Institutional Animal Care and Use Committee (IACUC) at the University of Colorado Anschutz Medical Campus under Protocol #00111 and performed in accordance with National Institutes of Health (NIH) guidelines. C57BL/6J animals were used for all experiments (RRID:IMSR_JAX:000664). Mice were housed separately by sex in groups of 1–5 mice/cage under a 14:10 light:dark cycle with controlled temperature and 35% humidity with *ad libitum* access to food (16% kcal fat) and water. Mice were euthanized by CO_2_ asphyxiation followed by cardiac puncture. Spleens were removed, then dissociated using a gentleMACS Octo Dissociator with Heaters (Miltenyi Biotec, cat. #130-096-427) using the preprogrammed 37C_m_SDK_1 protocol and the spleen dissociation kit (Miltenyi Biotec, cat. #130-095-926). T cells were isolated via negative selection with Dynabeads Untouched Mouse T Cells protocol (Invitrogen cat. # 11413D). Isolated mouse T cells were resuspended in culture medium: 1X RPMI Medium 1640 (Gibco, cat. # 11875093), 2 mM Glutamine, 10% FBS, 100 U/mL penicillin/streptomycin, 1X 2-Meracptoeethanol (Gibco, cat. # 21985023). Cells were plated at a concentration of 1.0 × 10^6^ per mL and activated in a 1:1 ratio with Dynabeads Mouse T-Activator CD3/CD28 (Gibco, cat. # 11452D) and 30 U/mL recombinant human interleukin-2 (Gibco, cat. # PHC0026) and allowed to culture for 48 hours. Cells were subsequently passaged at a concentration of 5.0 × 10^5^ cells per mL.

## METHOD DETAILS

### TSSG Stimulation and Inhibition

Fibroblasts or RPE cells were seeded at 4.0×10^5^ cells per 10-cm plate. Following overnight culture, cells were then transfected with exogenous nucleic acid or treated with 5 ng/mL TNFα (R&D Systems, cat# 210-TA-100). All nucleic acids (100 bp dsDNA at 250 ng/mL,^83^ 100 bp ssDNA at 500 ng/mL,^83^ ssRNA at 62.5 ng/mL (New England BioLabs, cat# N0364S), and dsRNA mimetic pIC (InVivoGen, cat# tlrl-picw) at 31.25 ng/mL) were transfected with Lipofectamine3000 (Invitrogen) at a ratio of 1 µg nucleic acid:2 µg of Lipofectamine3000 (1 mg/mL stock). All experiments were harvested 24 hours following transfection or treatment unless otherwise noted. The JAK1/2 inhibitor ruxolitinib (Selleck, cat# S1378) (dissolved in DMSO) was used at a final concentration of 5 µM, added to overnight culture media. The IKKβ inhibitor TPCA1 (Selleck, cat# S2824) (dissolved in DMSO) was used at a final concentration of 2 µM, added to overnight culture media. The p300/CBP bromodomain inhibitor SGC-CBP-30 (Selleck, cat# S7256) (dissolved in DMSO) was used at a final concentration of 2 µM, added to overnight culture media. All inhibitors were added to the cell culture 1 hour before transfecting nucleic acid or treating with TNFα. The media was not replaced prior to transfection.

### Cell harvesting and fixing for DNA FISH

Following tissue culture experimentation, conditioned media was collected in individual 15 mL tubes, then cells were washed once in 1x phosphate buffered saline (PBS) and lifted with 0.25% Trypsin-EDTA. Trypsin was then quenched with the conditioned media from each plate, respectively. Cells were pelleted at 300 g for 3 minutes, then washed once with 1x PBS. Cells were pelleted again at 300 g for 3 minutes. The PBS wash was then aspirated, and the cell pellets were then resuspended in 2 mL 0.075M KCl (hypotonic solution, 5.6 g KCl/L distilled water) and placed in 37°C water bath for 15 minutes with gentle intermittent agitation to prevent settling. 1 mL of fixative buffer (3:1 methanol:acetic acid) was then added, mixed, and cells were centrifuged. The supernatant was then aspirated, and the pellet was re-suspended in 2 mL of fixative buffer. The process of pelleting and re-suspending in fixative buffer was repeated 2 more times. Finally, cells were re-suspended in 1

### DNA FISH

Fixed cells from the -20°C suspended in fixative buffer were added (200-300 µL, empirically determined) to each well of Lab-Tek II Chamber slides (Thermo Fisher Scientific, cat# 154534) and attached to glass via centrifugation at 270 g for 3 minutes. The fixative buffer was then aspirated from chamber wells, and chamber walls are then removed. The slides are then air dried for 5-10 minutes at room temperature. The slides were washed in a series of coplin jars as follows; 2 minutes in 2X SSC buffer (Promega, cat# V4261), 2 minutes in 70% (v/v) EtOH, 2 minutes in 80% (v/v) EtOH, and finally 2 minutes in 95% (v/v) EtOH. The slide was kept on the bench at room temperature for 5-10 minutes to air dry. Following drying, 4 µL of DNA FISH probe master mix was spotted onto the center of each well, then gently applying a glass cover slip (VWR, cat# 16004-314). FISH probe master mix was prepared as follows. For each individual well of a Lab-Tek II Chamber slide, 3.4 µL of hybridization buffer (Empire Genomics, cat # NC3252634), 0.3 µL of target probe **Table S8**, and 0.3 µL of control probe were mixed, and placed in a 78°C heat block for 5 minutes. The master mix was then vortexed and briefly centrifuged prior to spotting onto the Lab-Tek II Chamber slide. Rubber cement was then used to seal the cover slip onto the slide, preventing overnight desiccation of the DNA FISH probes. The glass slide was denatured on a heating block at 78°C for 4 minutes. The slide was then placed at 37°C overnight in a sealed humidified bag. The following day, rubber cement was removed from the edges of the cover slip, and the slide was washed in a series of coplin jars as follows; 2 minutes in 0.4X SSC pre-heated to 68°C, 2 minutes in 2X SSC with 0.05% (v/v) Tween-20 at room temperature, 5 minutes DAPI staining solution (PBS containing 10 g/L Bovine Serum Albumin (Research Products International, cat# A30075-250) and DAPI (EMD Millipore, cat# 268298-10) at 1:10,000), then 5 minutes in 1x PBS. Following the 1x PBS wash, glass cover slips were mounted without air drying, using ProLong Gold antifade reagent (Thermo Fisher Scientific, cat# P36934). Slides were then placed in 37°C incubator for 1 hour, followed by overnight curing in the dark at room temperature. Slides were imaged on a Zeiss Axio Imager M2 at multiple positions for each sample. Images were captured in 10 µm Z-stacks in 20 slices using Zeiss Zen 2 Pro. Each nucleus was assigned a number to track scoring. For each nucleus the number of green and red foci was assessed by hand. A nucleus was considered to have a TSSG if it had three copies of *IFNR* probe and two copies of control probe (3:2) or five *IFNR* and three control probe (5:4). At least 110 nuclei were counted for each replicate.

### Immunoblotting

Sample preparation, quantitation, and immunoblotting were carried out as previously described.^60^ Detection was completed by chemiluminescence using SuperSignal West Pico PLUS (Thermo Fisher Scientific cat #34580), and images were captured with an ImageQuant800 digital camera system (Amersham). The antibodies used are described in **Table S11**.

### siRNA Knockdowns

2 × 10^5^ fibroblast cells were plated in 10-cm plates and allowed to culture overnight. Next day, 5 μL of 20 μM stock siRNAs (**Table S9**) were complexed with 15 μL of lipofectamine 3000 in 600 μL of OptiMEM (Gibco, cat# 31985-070) and added dropwise to a 10 mL overnight culture at a final concentration of 10 nM. 48 hours later, pIC (7.8 ng/mL) was transfected or TNFα (5 ng/mL) was added to media to stimulate the *IFNR* TSSG.

### CUT&RUN

4 × 10^5^ fibroblast cells were plated on 10-cm plates and allowed to culture overnight. Next day, cells were pretreated with 2 µM TPCA1 for 1 hour prior to stimulation. pIC was transfected at a final concentration of 31.25 ng/mL, or TNFα was added at a final concentration of 5 ng/mL to the overnight culture. pIC transfection was performed by complexing pIC with Lipofectamine3000 in 600 μL OptiMEM at a ratio of 1 μg pIC to 2 μg lipofectamine (1 mg/mL), scaled accordingly per experiment. Following a 24-hour pIC transfection or 24-hour TNFα treatment, the cells were washed once with PBS and lifted with 0.25% Trypsin-EDTA (Thermo Fisher Scientific). Cells were pelleted at 300 g and washed once with PBS. Nuclei were harvested via EpiCypher CUTANA CUT&RUN protocol V2.0. In brief, nuclei were harvested by resuspension in 1 mL of nuclear extraction buffer (20 mM HEPES pH 7.9, 10 mM KCl, 0.1% (v/v) Triton X-100, 20% (v/v) Glycerol, 1x cOmplete Mini-Tablet (Roche, cat # 50-100-3270), 0.5 mM spermidine (Sigma, cat # S2501)) per 5 × 10^6^ cells. Nuclei were incubated on ice for 10 minutes, then pelleted at 600 g for 5 minutes. The supernatant was aspirated, and the nuclei were resuspended in 100 μL nuclear extraction buffer per 5 × 10^5^ nuclei. Nuclei were stored frozen at -80°C in aliquots and thawed as needed for each CUT&RUN. Each CUT&RUN was performed on 5.0×10^5^ nuclei from two independent biological replicates following the EpiCypher CUTANA CUT&RUN protocol V2.0. Sequencing library preparation was carried out using an NEBNext Ultra II kit per manufacturer’s instructions

### RNP-Editing

1.0 × 10^5^ fibroblast cells were seeded into one well of a 6-well plate and allowed to attach overnight. RNP transfections were performed using the crRNA:tracrRNA duplex following the manufacturer’s recommended protocol (IDT). In brief, individual crRNAs (**Table S12**) were annealed to tracrRNA in equal molar amounts (1 μM each) using a thermal cycling program starting at 95°C and cooling to 12°C with 30 second holds at each temperature. Duplexed guide RNAs were then complexed with Cas9 protein in OptiMEM (Thermo Fisher Scientific) and packaged in Lipofectamine3000 (Thermo Fisher Scientific). The transfection mix was added dropwise to overnight cell cultures and allowed to incubate for 16 hours. The next day, transfection media was aspirated and replaced with fresh media. Cells were expanded into 10 cm^2^ plates prior to diluting 350 cells into 100 mL (0.35 cells per 100 μL) and plating these cells across 10 96-well plates. Media was replaced every 72 hours until wells were ready to split. Wells were then replicated into a DNA screening well and master plate well. For each well, the media was aspirated washed with 100 μL PBS, then trypsinized with 30 μL 0.25% Trypsin-EDTA. The well was quenched by the addition of 80 μL media, pipetted up and down to achieve single cell suspension, then 80 μL was added to the DNA plate, while the remaining volume was added to a master plate. The following day, DNA was isolated from the DNA screening plate. Media was aspirated and each well was washed with 100 μL of PBS, and the plate was briefly drained. 50 μL of lysis buffer (10 mM Tris-HCL pH 7.5, 10 mM EDTA, 10 mM Sodium Chloride, 0.5% (v/v) Sarkosyl) with Proteinase K (100 ng/mL) was added to each well and incubated overnight at 37°C. A 96-well rubber lid seal was used for each plate to collect evaporation, which was centrifuged back into each corresponding well the following morning. Salt precipitation of DNA was then performed by adding 100 μL of 0.075 mM sodium chloride in ice cold 100% (v/v) ethanol to each well and incubating for 20 min on ice. Precipitated DNA was then pelleted by centrifugation at 2000g for 15 min at 4°C. The wells were then washed once with 150 μL of ice cold 70% (v/v) ethanol, then allowed to air dry at RT for 15 min. DNA was then resuspended in 60 μL of water. For subsequent PCR screening, 2 μL of DNA was used as template in a 50 μL PCR reaction.

### Cas9-EZH2 Tethering

EZH2 [FL]-dCas9 (Addgene, cat # 100086), dCas9 (Addgene, cat # 100091) and gRNA_cloning vector (Addgene, cat # 41824) plasmids were purchased directly from Addgene and verified by sequencing. sgRNA plasmids were constructed by cloning of the sgRNAs into the gRNA_cloning vector (**Table S10**). A HiSpeed Plasmid Maxi Kit (QIAGEN, cat # 12663) was used to purify the plasmids for use in transfection. Verification of all Addgene plasmids was performed by full plasmid sequencing from Genewiz. To design gRNAs flanking the *IFNR* RelA site, ∼750 bp region was input to the Custom Alt-R CRISPR-Cas9 guide RNA tool from IDT. The gRNAs were chosen based on having a relatively low off-target risk, while having relatively high on-target potential. Each gRNA was incorporated into two 60-mer oligonucleotides as forward and reverse inserts. The two 60 bp oligonucleotides harbor overlap sequences and the sequences for U6 promoter and tracRNA in forward and reverse inserts respectively, as previously described ^55^. The 80 bp forward and reverse oligonucleotides were purchased from Sigma-Aldrich. The forward and reverse oligonucleotides were extended and annealed for each gRNA in a reaction containing 4 µL of 5X Phusion HF buffer, 0.4 µL of 10 mM dNTPs, 1 µL of forward oligo, 1 µL of reverse oligo, 0.2 µL of Phusion polymerase (Thermo Fisher Scientific, cat # M0530) and 13.4 µL of nuclease free H20 and using 3 cycles of extension using parameters as 94°C for 7 minutes, 94°C for 30 seconds, 60°C for 30 seconds, 72°C for 30 seconds and 72°C for 5 minutes. Immediately after completion of PCR, the tubes were placed on the bench to cool until room temperature. The 100 bp annealed amplicons were gel purified from 2% agarose (w/v) using after electrophoresis and cloned into gRNA_cloning vector. The gRNA_cloning vector (Addgene, cat # 41824) was linearized via restriction digestion using *Afl*II enzyme. The linearized vector was separated in 0.6 % (w/v) agarose gel electrophoresis and purified using QIAquick gel extraction kit (Qiagen, cat # 28704). Each sgRNA amplicon was cloned into gRNA_cloning vector via Gibson assembly using the Gibson assembly master mix (NEB) in a reaction containing 3:1 molar ratio of gRNA amplicon to gRNA_cloning vector at 50°C for 30 minutes. Afterwards, 2 µL of the assembly reaction mixture was transformed via heat shock into the C3040H competent cells (NEB, cat # C3040H). 200 µL of LB media was added to transformed cells and incubated at 37°C with vigorous shaking for 1 hour. 20 µL of the transformed culture was spread on Ampicillin LB plate and allowed to incubate at 37°C overnight. Colonies with correct plasmids containing the sgRNA inserts in the gRNA_cloning vector were confirmed by Sanger sequencing. 1 × 10^6^ fibroblast cells in DMEM were seeded for overnight culture in 10 cm plates. Transfection of the dCas9-EZH2 plasmids and sgRNA containing plasmids was performed on the next day. 30 μL of Lipofectamine 3000 (Invitrogen, cat # L3000001) was added in 270 μL of OptiMem (Gibco, cat # 31985070) in an Eppendorf tube (tube A) and allowed to sit for 5 minutes after mixing. Similarly, 18 μL of P3000 reagent, 3 µg of plasmid DNA (1.5 µg of dCas9 and 1.5 µg combined of the 6 sgRNAs) and the remaining volume of OptiMem were mixed to a total volume of 300 µL in another Eppendorf tube (tube B) and allowed to sit for 5 minutes. The contents from tube B were added and mixed into the contents of tube A and allowed to sit for 15 minutes. After the incubation time, the 600 µL mixture was slowly added to the fibroblasts in the 10-cm plate in a dropwise manner. The media was replaced with fresh DMEM after overnight transfection. 24 hours following transfection, 5 ng/mL TNFα was added to the cultures for 24 hours to induce *IFNR* TSSG.

### Mouse T Cell Experimental Conditions

T cells were plated at a concentration of 1.0 × 10^6^ cells per mL and treated with 31.25 ng/mL pIC complexed in Opti-Mem and Lipofectamine3000, 5 ng/mL recombinant mouse TNFα (Invitrogen, cat. # RMTNFAI), or associated vehicle and allowed to incubate for 24 hours. Cells were collected in culture media. Protein lysates and fixed cells for FISH applications were processed as described above. Copy-number was assessed via FISH with a mouse Chr16 *IFNR* probe (Empire Genomics, RP24-335D21) and mouse Chr16 control probe (Empire Genomics RP24-361P17).

### Transcriptome library preparation and sequencing

Total RNA was harvested using the AllPrep Mini Kit (Qiagen, cat # 80004) with the optional on-column DNase treatment (Qiagen, cat # 79254) per manufacturers recommendation. RNA quality was assessed using an Agilent 2200 TapeStation and quantified by Qubit (Life Technologies). Poly(A)+ RNA enrichment, and strand-specific library preparation were carried out using a Universal Plus mRNA-Seq with NuQuant (Nugen/Tecan, part # 0520). Paired-end 150 bp sequencing was carried out on an Illumina NovaSeq 6000 instrument by the Genomics Shared Resource at the University of Colorado Anschutz Medical Campus.

## QUANTIFICATION AND STATISTICAL ANALYSIS

Data preprocessing, statistical analysis, and plot generation for all datasets were carried out using R (R 4.0.1/RStudio 2022.02.0/Bioconductor 3.11) as detailed in supplementary materials. Differentially expressed genes from transcriptome experiments were identified using DESeq2 and peak calls for CUT&RUN were made using HOMER ^56,57^, see supplementary materials for details. All figures were assembled in Adobe Illustrator v25.4.5.

### Analysis of RNA-seq data

Sequencing data quality was assessed using FASTQC (v0.11.9) and FastQ Screen (v0.15.2).^84^ Trimming and filtering of low-quality reads was performed using bbduk from BBTools (v39.01) and fastq-mcf from ea-utils (v1.04). Alignment to the human reference genome (GRCh38) was carried out using HISAT2 (v2.1.2.1) in paired, spliced-alignment mode against a GRCh38 index and Gencode v33 basic annotation GTF, alignments were sorted and filtered for mapping quality (MAPQ > 10) using Samtools (v1.16.1),^85^ and duplicates marked using Picard (v3.0.0). Quality assessment of final mapped reads was conducted using RSeQC (v5.0.1). Gene-level count data were quantified using HTSeq-count (v2.0.2) with the following options (--stranded=reverse – minaqual=10 –type=exon --mode=intersection-nonempty) using a Gencode v33 GTF annotation file.^86^ Differential gene expression in stimulated vs control treatments were evaluated using DESeq2 (version 1.44.0),^87^ adjusting for batch effects, with q < 0.1 (10% FDR) as the threshold for differential expression.

### CUT&RUN Data processing

Sequencing data quality was assessed using FASTQC (v0.11.9) and FastQ Screen (v0.15.2). Trimming and filtering of low-quality reads was performed using bbduk from BBTools (v39.01) and fastq-mcf from ea-utils (v1.04). Alignment to the human reference genome (GRCh38) was carried out using BWA (v0.7.17, alignments were sorted and filtered for mapping quality (MAPQ > 10) using Samtools (v1.16.1), and duplicates marked using Picard (v3.0.0). Quality assessment of final mapped reads was conducted using RSeQC (v5.0.1). The Homer suite (version 4.11.1) was used for generation of bedGraph files for genome browser visualization with the Homer makeUCSCfile module (-ref hg38, -norm 1e6).^88^

### Peak Calling and Global binding profile of CUT&RUN

Downstream analysis was performed using HOMER.^88^ HOMER makeTagDirectory and findPeaks tools were employed to call peaks using a Poisson distribution to identify regions with statistically enriched binding for each target against a matching IgG control. Antibody-target specific findPeaks options were used for peak calling: RelA, ‘-style factor -center -F 3 -L 3’; epigenetic marks and MCM2, ‘-style histone -size 300 -minDist generated from bam files for heatmap visualizations using deepTools bamCoverage (v3.5.1), with normalization using counts per million (CPM) per bin with 50 bp bins. Comprehensive peak call lists for each target were generated by merging peak calls that overlapped by at least 1 bp across all conditions and replicates for a given experiment using the bedtools merge function (v2.30.0), followed by centering on the merged peak center. Heatmaps were generated using comprehensive peak call sets and associated Bigwig files with deepTools computeMatrix and plotHeatmap functions centered on the peak center with 50 bp bins, 5 kb windows, and sorted by the window mean on the treatment conditions.

### Identification of inducible binding of RelA and MCM2 from CUT&RUN

Inducible peaks were identified using the HOMER findPeaks tool for a given treatment compared to untreated control conditions for each antibody target with the same parameters as above. These peaks are referred to as inducible peaks herein. Peaks that overlapped by at least 1 bp with ENCODE blacklist regions were removed. Consensus inducible peaks were then generated by identifying peaks with at least one bp overlap across the two replicates. For instances of one-to-many relationships between replicates for histone marks and MCM2, the peak with the largest associated fold-change was retained for downstream analyses in that replicate. Consensus peak calls were then annotated with HOMER annotatepeaks.pl function with GRCH38 as the reference. Volcano plots, p-value vs p-value, and fold-change vs fold-change plots were generated from these consensus inducible peaks in R. Inducible MCM2 heatmaps were generated from the consensus inducible peaks using the same methodology described above.

### Identification of co-inducible RelA and MCM2 binding from CUT&RUN

The consensus inducible MCM2 peaks were filtered based on regions that overlapped with the TNFα inducible RelA peak call lists for both replicates, by at least one bp, generating the class one (inducible MCM2 only), and an inducible MCM2-RelA class of peaks. The inducible MCM2-RelA peaks were then sub-divided into class two (repetitive gene families), and class three (non-repetitive gene families) based on the Gene Duplication Database. Lastly, Gene-set Enrichment Analysis was performed for MCM2, TNFα vs control peaks. Replicates were run individually with their respective fold-changes. P-value, fold-change, and NES results were averaged across the two replicates.

## Supplementary Material

**Figure S1.**
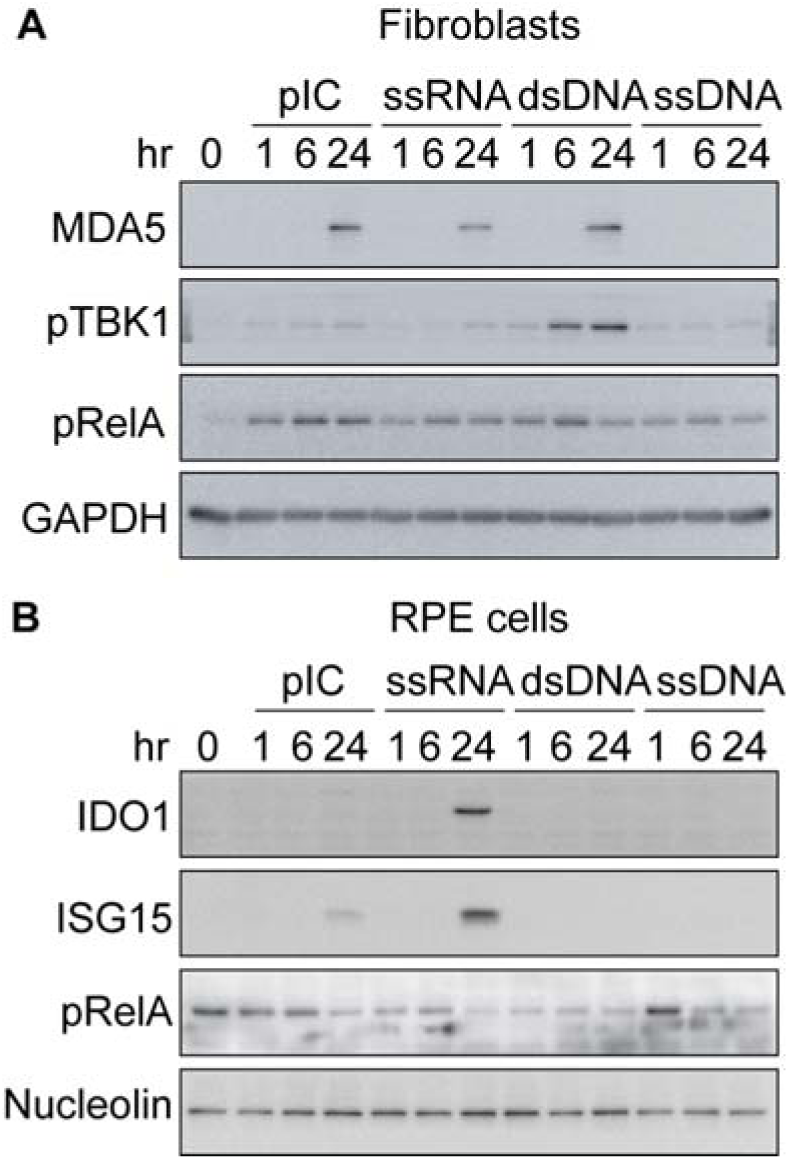
Innate immune activation by nucleic acid transfection. (**A**) Immunoblot characterization of a time course of nucleic acid treatments in fibroblasts. Time course experiment was completed in parallel with the 24-hour nucleic acid transfections shown in **Figure. 1D**. (**B**) Immunoblot characterization of a time course of nucleic acid treatments in Retinal Pigment Epithelial (RPE) cells. Time course experiment was completed in parallel with the 24-hour nucleic acid transfections as shown in **Figure. 1F**.

**Figure S2.**
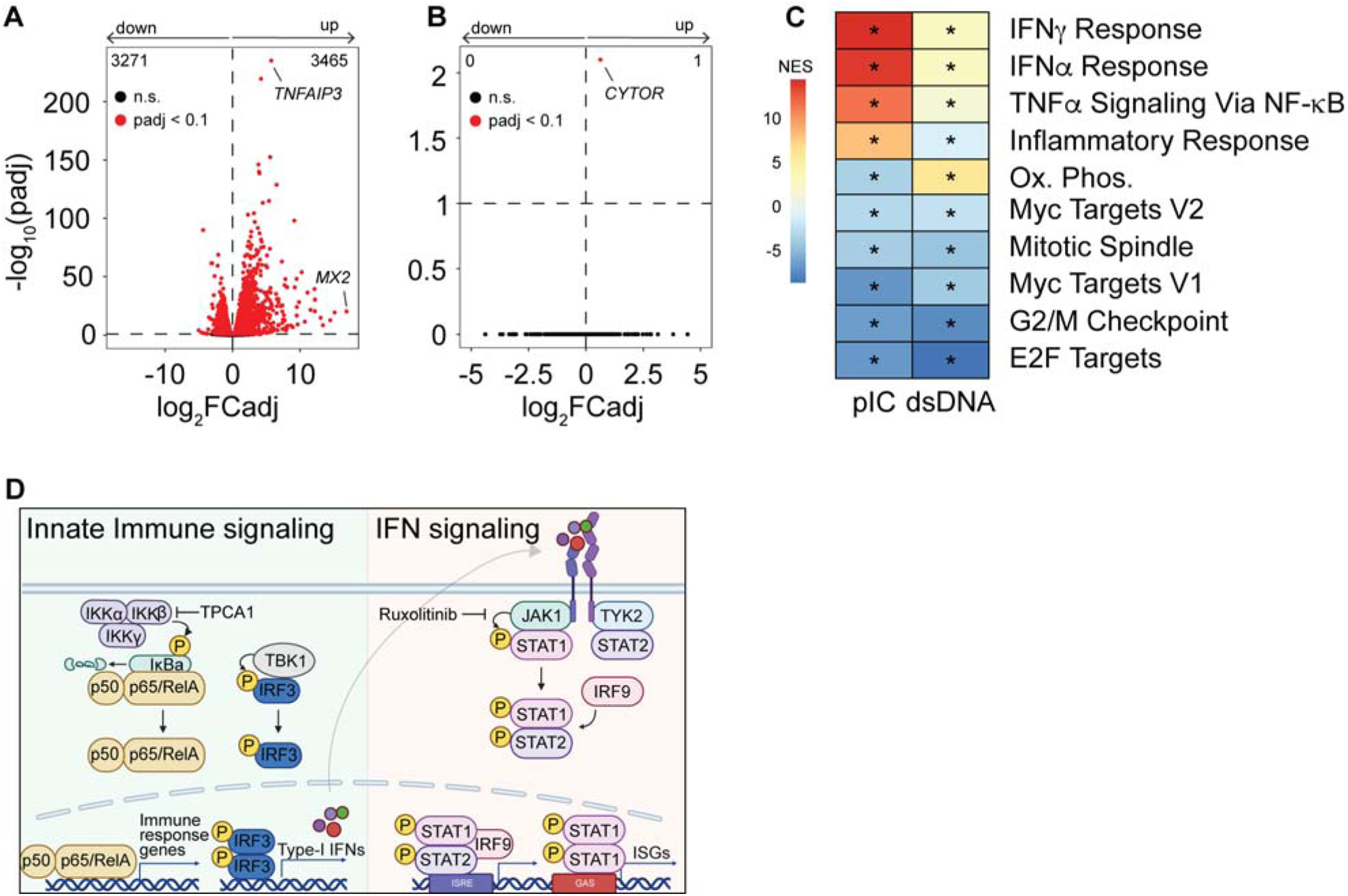
Pharmacological and genetic characterization of the regulation of the *IFNR* TSSG. (**A**) and (**B**) Volcano plots summarizing transcriptome changes (q<0.1) in RPE cells following 24-hour transfection of pIC (31.25 ng/mL, **A**) or dsDNA (250 ng/mL, **B**). (**C**) Heatmap of gene-set enrichment analysis (GSEA) of Hallmarks gene sets. Asterisks in the box denote significance (q-value < 0.1) after Benjamini-Hochberg adjustment of *p* values. (**D)** Schematic of innate immune signaling via the NFκB and IRF3 pathways, pharmacological inhibitors and targets used in Figures 2D-G and **2L-O** are noted. Created with BioRender.

**Figure S3.**
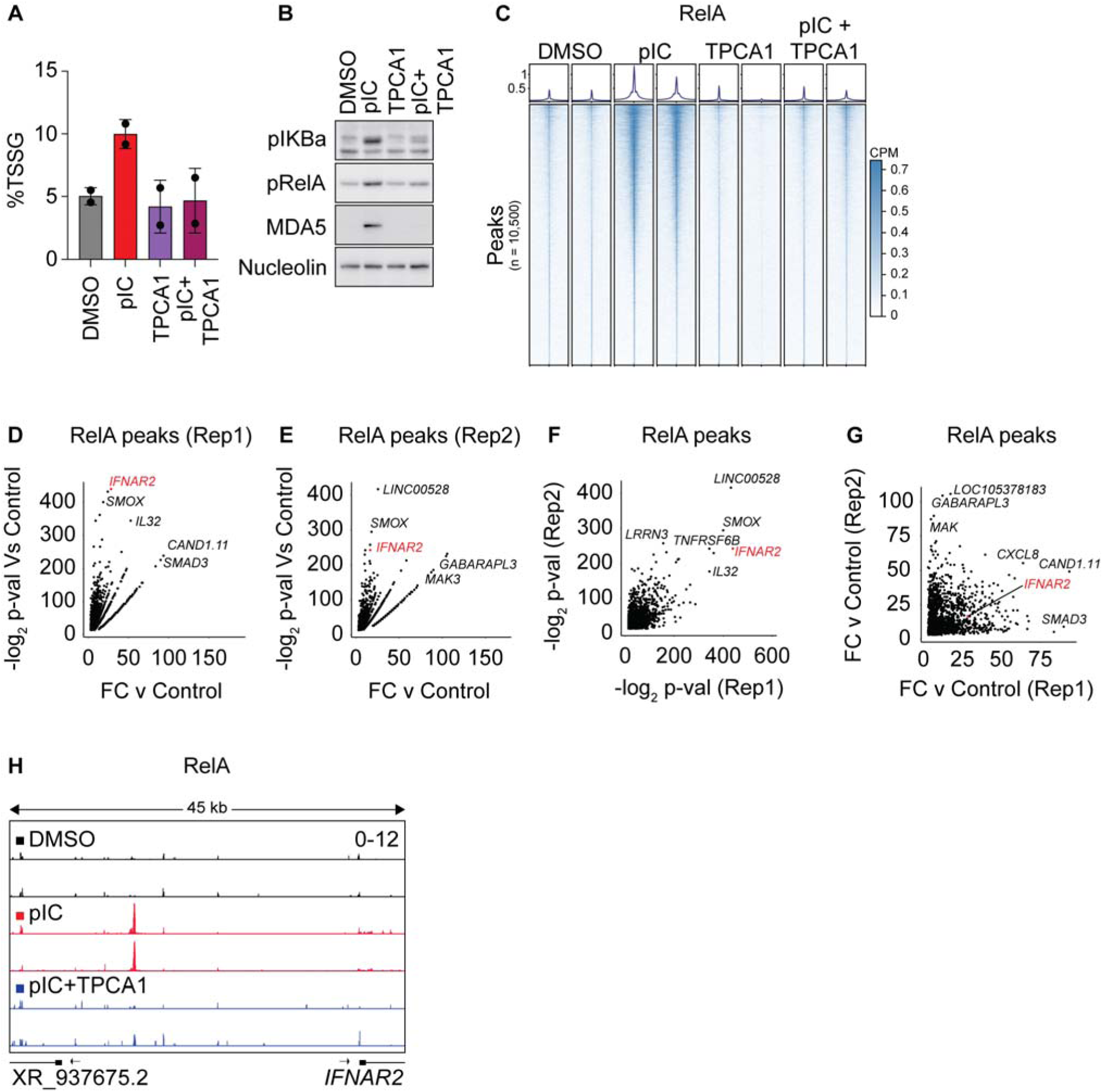
RelA binds a site upstream of *IFNR* locus. (**A**) Percentage of fibroblasts undergoing *IFNR* TSSG for two biological replicates utilized for all pIC transfected CUT&RUN experiments (n = 2). (**B**) Immunoblot analysis of RelA activation for CUT&RUN samples in **A**. (**C**) Heatmaps of RelA signal surrounding HOMER peak calls from two biological replicates of RelA CUT&RUN for the indicated treatment. Each row represents a peak that was called in at least one treatment condition against a matched IgG control. Overlapping peaks across treatment conditions were merged. Columns range from -2.5 kb to 2.5 kb from peak center with 50 bp bins. (**D-E**) Fold change vs -log_2_ p-value plot for RelA HOMER peak calls in pIC transfected cells for each replicate. (**F**) Scatter plot of -log_2_ p-values from peaks from two biological replicates of pIC-transfected RelA CUT&RUN. (**G**) Scatter plot of fold changes of peaks from two biological replicates of pIC-transfected RelA CUT&RUN experiments. (**H**) RelA CUT&RUN tracks from two biological replicates treated as indicated. Unless otherwise noted, statistical significance was assessed by a 2-tailed student’s t-test. Black dots represent independent biological replicates. Error bars indicate SD.

**Figure S4.**
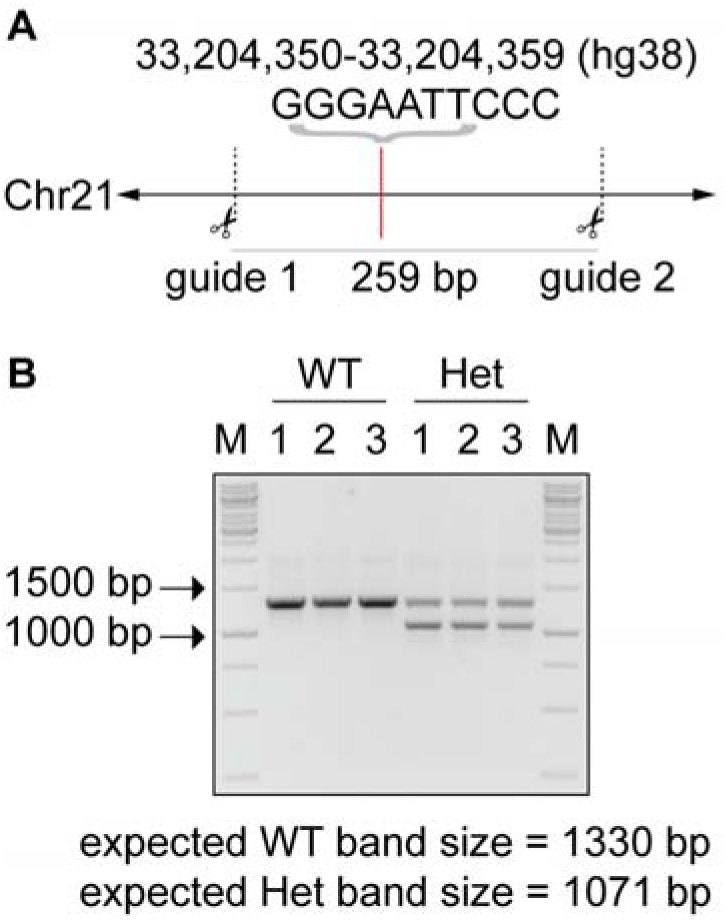
Epigenetic Control of *IFNR TSSG*. (**A**) Schematic of CRISPR-Cas9 gRNA cut sites indicating region deleted in Het-RNP edited lines. The location of the RelA binding site is indicated by the sequence of nucleotides and the red line. (**B**) Agarose gel of PCR from three clones retaining both WT alleles (WT-RNP) and three clones having a single RNP-edit (Het-RNP) which were pooled for subsequent experiments.

**Figure S5.**
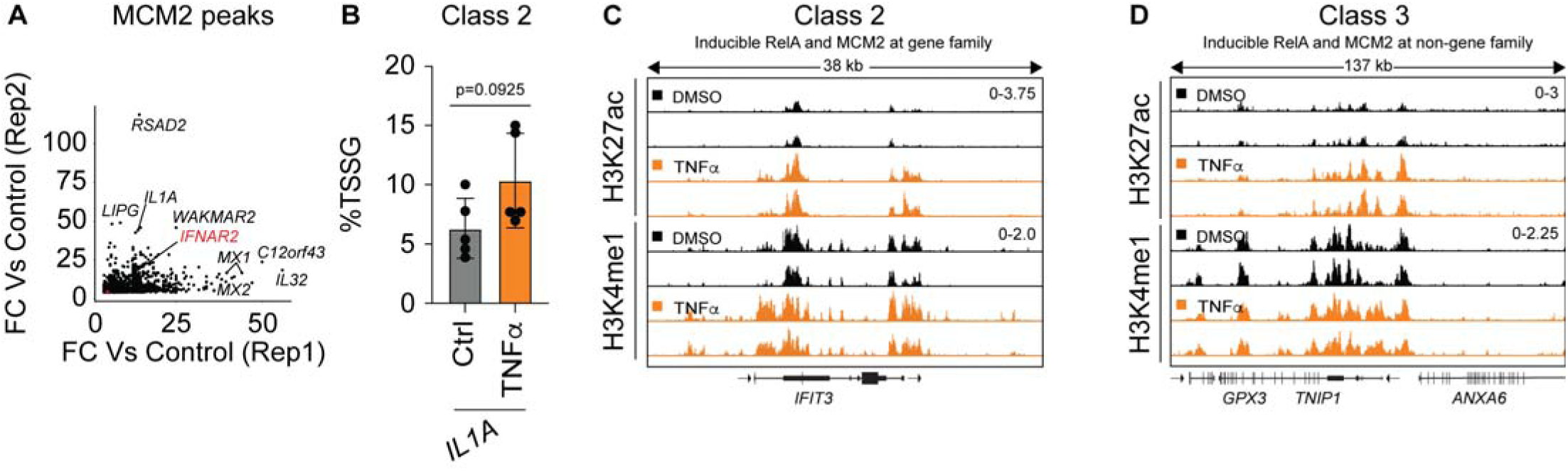
MCM2 and RelA colocalization control a TSSG program induced by innate immune signaling. (**A**) Scatter plot representing fold change from inducible RelA-chromatin interactions from both biological replicates of TNFα RelA CUT&RUN experiments. (**B**) Percentage of cells undergoing *IFNR* TSSG in cells treated with TNFα (5 ng/mL) for 24 hours (n = 5). (**C**) H3K27ac and H3K4me1 CUT&RUN tracks from two biological replicates at the Class 2 gene *IFIT3*. (**D**) H3K27ac and H3K4me1 CUT&RUN tracks from two biological replicates at the Class-3 gene *TNIP1*. Unless otherwise noted, statistical significance was assessed by a 2-tailed student’s t-test. Black dots represent independent biological replicates. Error Bars indicate SD.

## Supplementary Table Legends

**Table S1. DNA FISH Summary Data.** Table S1 provides summary data for all DNA FISH experiments in this manuscript. Tabs are labelled according to figure panel.

**Table S2. DESeq2 Results.** Table S2 provides DESeq2 results for transcriptomic analysis of fibroblasts (**Figures 2A** and **2B**) and RPE cells transfected with pIC and dsDNA (**Figures S2A** and **S2B**).

**Table S3. RNAseq GSEA.** Table S3 provides GSEA Hallmark Analysis for pIC and dsDNA treated fibroblasts and RPE cells (**Figures 2C** and **S2C**).

**Table S4. CUT&RUN Homer peak calls for transcription factors, MCM2, and epigenetic marks versus IgG.** Table S4 provides Homer peak calls for RelA and MCM2 CUT&RUN experiments versus IgG (**Figures 3O**, **S3C,** and **S6A)**.

**Table S5. CUT&RUN Homer peak calls for transcription factors, MCM2, and epigenetic marks versus control.** Table S5. Provides Homer peak calls for RelA, MCM2, and epigenetic marks (H3K4me1, H3K4me3, H3K27ac1, H3K36me3, H4K20me1, K3K37me1, K3K27me3, H3K9me1, H3K9me3), for cells treated with pIC or TNFL versus control treatment (0.05% DMSO or PBS+0.1% BSA). (**Figures 3P – 3S, S3D – S3G, 4N, 4P, 5B, 5C, 5E 5F, 5H, 5I, 6B, 6E,** and **S6A**)

**Table S6. MCM2 CUT&RUN GSEA.** Table S6 provides GSEA Hallmark analysis of Homer peak calls for MCM2 CUT&RUN for cells treated with TNFL versus control treatment (PBS+0.1% BSA) (**Figure 6F**).

**Table S7. Class I, II, and III gene lists with CUT&RUN peak coordinates.** Table S7 provides Homer peak call coordinates for CUT&RUN data associated with Class I, II, and III genes (**Figure 6I – 6J** and **6O**).

**Table S8. FISH Probe IDs.** Table S8 provides BAC clones used for FISH probe production by Empire Genomics for all FISH probes used in this study.

**Table S9. siRNA sequences.** Table S9 provides all Invitrogen Silencer Select siRNA sequences used in this study.

**Table S10. dCas9-tethering gRNAs.** Table S10 provides sequences for all gRNAs used for dCas9 tethering experiments in **Fig 4**.

**Table S11. Antibodies.** Table S11 provides information for all antibodies used in this study for both immunoblot and CUT&RUN.

**Table S12. Cas9-RNP gRNAs and primers.** Table S12 provides sequences for gRNAs used to edit the *IFNAR2* TCE as well as primers used for screening of edited clones (**Figures 5A** and **5B**).

